# Linking Tissue Morphology and Tissue Healing in a Cell-Fate Model

**DOI:** 10.1101/2024.08.25.609579

**Authors:** Somya Mani, Tsvi Tlusty

**Affiliations:** Institute for Basic Science-Center for Soft and Living Matter, South Korea; Konrad Lorenz Institute for Evolution and Cognition Research, Austria; Department of Physics, Ulsan National Institute of Science and Technology, South Korea

**Author notes:** Correspondence: Somya Mani.

**Keywords:** Cell-fate, cell differentiation, cell migration, cellular neighborhood, tissue domains, tissue healing

## Abstract

Multicellular tissues are immensely diverse, and yet are notably similar at the microanatomical level – most tissues are organized into distinct domains with well-defined cellular compositions and precise adjacencies among themselves. Concurrently, tissues across organisms are also similar in an important functional property: the ability to heal from injury, even in organisms with poor regenerative capacities, such as mammals. Both the cellular organization within tissues and the ability of tissues to heal are outcomes of developmental processes, suggesting that the two may be intrinsically linked. In this work, we explore this connection using an agent-based model of developmental cell-fate decisions. Our model produces a rich diversity of tissue morphologies: By tuning only four parameters, which control the density of intercellular interactions and the propensity of cellular differentiation, we generate tissue structures ranging from disordered and sparse to tissues organized into dense, contiguous domains. Only tissues with large contiguous domains had the ability to heal from injuries, thus demonstrating that tissue healing is strongly coupled to tissue morphology. Moreover, model-generated tissues predominantly heal through the replacement of injured cells by cells dividing in the neighborhood, which recapitulates natural mechanisms in animals as well as plants. More generally, our modeling framework allows a systematic sampling of diverse developmental programs and reproduces healing mechanisms in organisms as phylogenetically distant as plants and animals. Our work thus points to general patterns which often are difficult to directly compare in empirical research. Despite the multiple evolutionary origins of multicellularity, the demonstrated link between tissue domain-level organization and healing capacity may represent a unifying feature of multicellular life.

## INTRODUCTION

Tissues are systems of cells of one or more cell-types that together perform specific functions beyond what any single cell is capable of (Adler et al., 2023). Tissue level organization is posited to be essential for the evolution of functionally specialized cells in multicellular organisms (Pavlicev et al., 2024). Correspondingly, tissue level organization is widely distributed across the different multicellular lineages of animals (Technau and Steele, 2011), land plants (Oliva and Lister, 2023), brown algae (Theodorou and Charrier, 2021) and fungi (Nagy et al., 2023). In keeping with the diversity of these organisms, their tissue compositions are also diverse, ranging from the Cnidarian *Hydra* which contains two tissue layers that have 3-4 cell-types (Holstein, 2023), to complex animals like humans which contain at least sixty tissues that are composed of tens of cell-types (Hatton et al., 2023).

Irrespective of cellular complexity, cells within any tissue interact closely with one another using molecular signals and mechanical cues. These inter-cellular interactions shape cellular decisions to divide, die, differentiate or migrate (O’Connor et al., 2010). These cellular decisions in turn dictate tissue form and function (Ramos et al., 2024), and tissues across organisms tend to organize into distinct microanatomical domains that are characterized by their cellular compositions and form niches for inter-cellular interactions: For example, there are three distinct domains in the mammalian intestinal epithelium: the crypt base which contains the stem cells, the crypt walls which contain transit amplifying cells and the villi which contain differentiated cells (Clevers, 2013). Similarly, in plants, the shoot apical meristem is composed of three domains: the central zone which contains the stem cells, the organizing center whose cells provide hormonal cues to the central zone, and the peripheral zone which contains differentiated cells (Gaillochet et al., 2015). At the same time, cellular interactions are also essential for healing of tissues from injuries: For example, in mammalian muscles (Koike et al., 2022) and spinal cord (Brennan et al., 2022), and in plants (Sugimoto et al., 2019).

In this work, we developed a spatial agent-based model to explore how the control of cell-fate decisions by inter-cellular interactions regulates cellular organization within tissues, and how tissue healing is related to tissue cellular organization. Our model produces a rich variety of tissue-level organizations which, similarly to real tissues, are arranged into distinct domains. The model’s tissues vary widely in the relative arrangements of these domains, ranging from those with sparse cellular distribution to full and densely occupied grids, and tissues with small, disperse domains to large, contiguous domains. Crucially, we found tissue healing to be strongly linked with tissue form: almost all regenerative tissues were densely occupied with cells which formed large and contiguous domains. Moreover, the model’s tissues healed predominantly via a localized response where cells lost upon injury were replaced through cell-division and differentiation of neighboring cells. This is the most probable mechanism that emerges robustly across a wide sampling of cellular interactions, though specific organisms may employ alternative strategies shaped by their evolutionary histories. Our model also produces experimentally testable predictions since our parameters represent elementary developmental processes which are amenable to experimental manipulation. Specifically, we predict that increasing the level of inter-cellular interactions should cause tissues with contiguous domains to become disordered and reduce their ability to heal.

### MODEL OF CELLULAR INTERACTIONS AND CELL-FATES

A tissue in the model is composed of *N* distinct cell types. These cells are distributed across the tissue’s body: In the model, the tissue is housed in a 2-dimensional grid with *G*^2^ grid-squares and periodic boundaries (Fig.1(B)). Multiple cells are allowed to reside within any grid-square: *C*^*n,g*^ gives the number of cells of cell-type *n* in the grid-square *g*, for *n* ∈ {1, 2, 3, …*N*} and *g* ∈ {1, 2, 3, …*G*^2^}.

**Figure 1.**
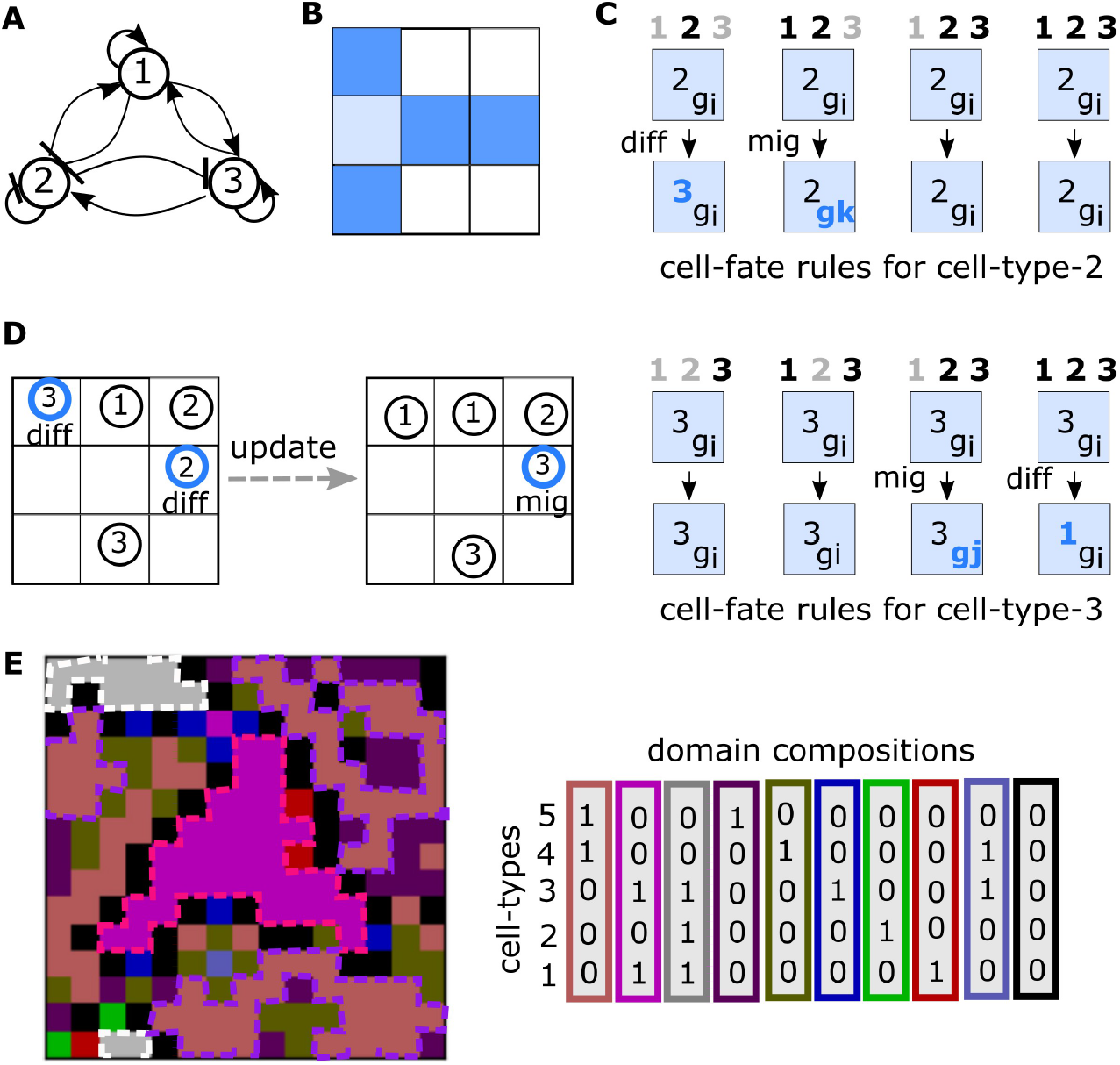
An example of cellular interactions and cell-fate rules for a tissue with *N* = 3 cell-types in a *G* = 3 grid. **(A)** Cellular interactions: Circles represent distinct cell-types. Sharp arrows indicate stabilizing interactions and blunt arrows indicate destabilizing interactions. Cell-types *1* and *3* are intrinsically stable whereas cell-type *2* is intrinsically unstable. **(B)** Adjacent grid-squares: The grid has periodic boundaries and the dark blue squares are all adjacent to the light blue square. **(C)** An example of Cell-fate rules for cell-types that interact according to (A): Cell-type-1 is intrinsically stable and has only stabilizing interactions, therefore, it neither differentiates not migrates in any cellular neighborhood. We show here cell-fate rules for cell-type-2 (top) and cell-type-3 (bottom). Light blue squares in the first row represent the grid-square occupied by the cell of interest: The cell-type and the location of the grid-square are indicated within the square. The numbers in bold written above the square represent the cellular neighborhood: black numbers indicate the presence of the corresponding cell-type and grey numbers indicate the absence of the corresponding cell-types. Arrows point to the cell-fate of the cell of interest in the corresponding cellular neighborhood: Cellular decisions of unstable cells to differentiate or migrate are are highlighted in blue font.**(D)** A tissue represented by the grid on the left updates to the grid on the right following Cell-Fate Rules given in (C). **(E)** An example of a stable *N* = 5, *G* = 15 tissue. Colors of grid-squares indicate their cellular compositions. We call contiguous regions of the tissue that have the same composition a *domain*: Cells in the center of the domain have a cellular neighborhood identical to the domain composition, while cells at domain edges have richer cellular neighborhoods. Domains have variable sizes: we highlight with dashed borders the three large domains in this tissue. The cellular compositions of these three domains are given alongside the image: 1 indicates the presence of at least one cell of the corresponding cell-type while 0 indicates absence of the corresponding cell-type. Black grid-squares are empty.

The geometry of the body is given by a distance matrix D: *D*(*g*_*a*_, *g*_*b*_) = *d* ∈ 𝕨 implies the grid-squares *g*_*a*_ and *g*_*b*_ are a Manhattan distance *d* apart. Specifically, *g*_*a*_ and *g*_*b*_ are *adjacent* if *D*(*g*_*a*_, *g*_*a*_) ≤ 1. In a 2-dimensional grid, each grid-square is adjacent to five grid-squares including itself (Fig.1(B)).

Cells in adjacent grid-squares are called *neighbors*. The *neighborhood* of any cell in grid-square *g* is given by a *N*-length binary vector *H*_*g*_: *H*_*g*_(*n*) = 1 if a cell of cell-type *n* is in the neighborhood of square *g*; *H*_*g*_(*n*) = 0 otherwise. There are 2^*N*^ distinct possible neighborhoods, which can be arranged in the standard binary ordering: 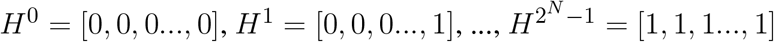.

By allowing multiple cells to reside within the same grid-square, our model presents a coarser picture of cellular arrangements within tissues compared to standard models where the location, geometry and neighborhood of each cell (such as in cellular automaton (Lee et al., 1995) and vertex models (Farhadifar et al., 2007)) or even subparts of cells (such as in the cellular Potts model (Marée et al., 2007)) are explicitly described. Thus our formalism offers a more general representation of tissues since we do not impose constraints on the geometries or adjacencies of individual cells (Osborne et al., 2017). At the same time, the simplicity of our model as compared to the cellular Potts model and lattice-free models (such as in (Bock et al., 2010; Sandersius and Newman, 2008)) makes it computationally feasible to sample a broad space of tissue patterns.

### Tissue developmental rules

Cellular interactions are prescribed by the interaction matrix *I* (Fig.1(A), Methods: Generating tissue developmental rules): A cell of type *n*_*i*_ in square *g*_*a*_ interacts with a cell of type *n*_*j*_ in square *g*_*b*_ if *I*(*n*_*i*_, *n*_*j*_) ≠ 0 and *g*_*a*_ and *g*_*b*_ are adjacent. That is, cellular interactions in the model are short to medium-range and a cell can interact with another cell some tens of cells away. We do not impose any constraints on the form of cellular interaction and do not distinguish between molecular and mechanical signals (Pavlicev and Wagner, 2023). The model’s cellular interactions can represent short-range molecular signaling, which are known to be important during tissue development (Armingol et al., 2024), for example, during embryonic development across animals (Perrimon et al., 2012) and during the growth and development of tissues in the plant shoot (Gaillochet et al., 2015). The model’s cellular interactions can also represent mechanical signals (Chan et al., 2017) such as via the local modification of the extracellular matrix by one of the cells which impacts the behavior of nearby cells. For example, during skull development in mice, differentiated osteoblasts secrete collagen into the extracellular matrix and induce differentiation of nearby mesenchymal cells (Dang et al., 2023).

We call a cellular interaction *stabilizing* if *I*(*n*_*i*_, *n*_*j*_) = 1 and *destabilizing* if *I*(*n*_*i*_, *n*_*j*_) = −1. Particularly, a cell-type *n*_*i*_ is *intrinsically stable* if *I*(*n*_*i*_, *n*_*i*_) = 0 or 1 and *intrinsically unstable* if *I*(*n*_*i*_, *n*_*i*_) = −1. Intrinsically stable cells are destabilized in the presence of any destabilizing interactions and intrinsically unstable cells are stabilized in the presence of any stabilizing interactions. For example, the germline stem cells (GSC) in the worm *Caenorhabditis elegans* are intrinsically unstable and tend to differentiate into germ cells. Interaction of GSCs with distal tip cells stabilizes them by repressing their differentiation (Byrd and Kimble, 2009).

A stable cell maintains its cell-type and its position within the body, whereas an unstable cell either differentiates into a new cell-type or migrates to a new grid-square. In the model, we allow a migrating cell to move a randomly chosen distance *m* ≤ *G* away. In order to maintain tissue connectedness, cells in the model can only migrate through grid-squares that are already occupied, though they are allowed to migrate into an empty grid-square which is adjacent to an occupied square.

Additionally, unstable cells of the same cell-type behave differently in different cellular neighborhoods (Fig.1(C)): For example, let a cell of cell-type *n*_*i*_ be unstable in two distinct neighborhoods *H*^*a*^ and *H*^*b*^. In neighborhood *H*^*a*^, the cell differentiates into cell-type *n*_*j*_; i.e. *n*_*i*_|*H*^*a*^ → [*n*_*j*_, *m* = 0], while in neighborhood *H*^*b*^, the cell migrates; i.e. *n*_*i*_|*H*^*b*^ → [*n*_*i*_, *m* = *m*_1_]. In nature, the effect of such positional inputs on cell-fate are especially apparent in migratory multipotent stem cells, such as neural crest cells in zebrafish (Subkhankulova et al., 2023).

While for the most part, plasticity of cell-fate is restricted to stem cells, under special circumstances, such as stem cell ablation, differentiated cells can revert to a stem-like state capable of proliferation and differentiation (Yao and Wang, 2020). Hence in the model, we do not impose any restrictions on cell-fate potential, and instead allow hierarchical structures in cell-fate potential to emerge from the interplay of cellular interactions. Cell Fate Rules in the model comprise the set of all cell-fate transitions *n*_*i*_|*H*^*a*^ → [*n*_*j*_, *m*], such that *n*_*i*_, *n*_*j*_ ∈ {1, 2, …, *N*}, *m* ≤ *G* and *a* ∈ {0, 1, …, 2^*N*^ − 1} (see Methods: Generating tissue developmental rules).

In addition to cellular interactions, differentiation and migration, cells in the model can also divide and die (Methods:Updating tissue states). In nature, cell proliferation is often regulated by cell density (Fan and Meyer, 2021; Pavel et al., 2018). To capture this in the model, cell division and cell death are regulated by the number of cells in grid-squares: Let *TC*^*g*^ be the total number of cells in some grid-square *g*. Any cell in grid-square *g* divides into an identical cell-type with probability *P*_*div*_ = *max*(0, (10 − *TC*^*g*^)*/*9) and a cell dies with probability *P*_*death*_ = *min*(0.8, 0.02(*TC*^*g*^ − 10)). The fixed cell-division and death parameters we use lead to grid-squares having on average 10 cells; therefore the neighborhood of a single cell, which comprises five adjacent grid-squares (Fig.1(B)), is composed of 50 cells. The fixed parameters in these formulae can be tuned to suit different cellular densities.

The daughter cell of a dividing cell in grid-square *g* can be born either in the same square or in some adjacent square. In nature, many cells divide asymmetrically during embryonic development, and in adults, tissue stem cells divide asymmetrically to produce daughter cells with distinct fates (Sunchu and Cabernard, 2020). In the model, daughter cells are identical to each other immediately after cell division, but cells born in adjacent grid-squares can end up with distinct cell-fates, which allows us to capture the effects of asymmetric cell-division.

### Model parameters

We draw the interaction matrix *I* using parameters 0 ≤ *P*_*den*_ ≤ 1 and 0 ≤ *P*_*stable*_ ≤ 1: *P*_*den*_ controls the level of inter-cellular signaling – *I*(*n*_*i*_, *n*_*j*_) ≠ 0 with probability *P*_*den*_; *P*_*stable*_ controls cell-fate potential – *I*(*n*_*i*_, *n*_*j*_) is positive with probability *P*_*stable*_.

We draw Cell Fate Rules using parameter 0 ≤ *P*_*diff*_ ≤ 1: for some cell of cell-type *n*_*i*_ that is unstable in neighborhood *H*^*a*^, the cell differentiates with probability *P*_*diff*_ and migrates otherwise. In neighborhood *H*^*a*^, if cell-type *n*_*i*_ differentiates, the final cell-type *n*_*j*_ is chosen uniform randomly from among the N distinct cell-types. Instead if cell-type *n*_*i*_ migrates, the number of grid-squares it migrates is drawn uniformly from within a distance *m* ≤ *G*. Together, the interaction matrix *I* and the Cell Fate Rules comprise *tissue development rules* (Methods:Generating tissue developmental rules). *Tissue development rules*, once drawn, remain fixed throughout a simulation.

Finally, the parameter 0 ≤ *F*_*adj*_ ≤ 1 controls the location of daughter cells after cell division: A fraction 1 − *F*_*adj*_ of daughter cells are born within the same grid-square as the mother cell, and a fraction *F*_*adj*_ are born in a different adjacent grid-square. *F*_*adj*_ is a complex parameter that captures aspects of cellular organization within grid-squares, including cell shape (McKinley et al., 2018) and the directions of the axes of cell-division (Gillies and Cabernard, 2011).

At any time-step *t*, the state of the tissue is completely characterized by the cellular compositions of its grid-squares 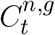 for all *n* ∈ {1, 2, …*N*} and *g* ∈ {1, 2, …*G*^2^}. At *t* = 0, we initialize the tissue by randomly placing 20 cells of random cell-types across the connected set of grid-squares [0, 1, 2, 3, 4]. We update the tissue by synchronously updating the state of each cell in the tissue according to the *tissue development rules* (Fig.1(D), Methods:Updating tissue states). We simulate tissue development in the model for *T* = 500 time-steps. We explore the spatial features and cellular compositions of tissues in the final time-step (Fig.1(E)), and analyze how tissues grow and heal from injuries.

## RESULTS

### Model-generated tissues are rich in cell-types

We first analyse how Cell Fate Rules regulate the cellular compositions of model-generated tissues. A majority of model-generated tissues across all parameters contain all 5 cell-types in the final time-step (Fig.2(A), Supplementary Information:FigS2). We find that the number of tissue cell-types increases with the probability of stabilizing interactions, *P*_*stable*_ (Pearson’s *r* = 0.15) and decreases with the propensity of cells to differentiate, *P*_*diff*_ (*r* = -0.41). Intuitively, we expect that increasing the number of cellular interaction partners should increase the probability of destabilizing interactions. In keeping with this, we find that the number of tissue cell-types decreases with *P*_*den*_ which controls the probability of inter-cellular interactions (*r* = -0.14).

**Figure 2.**
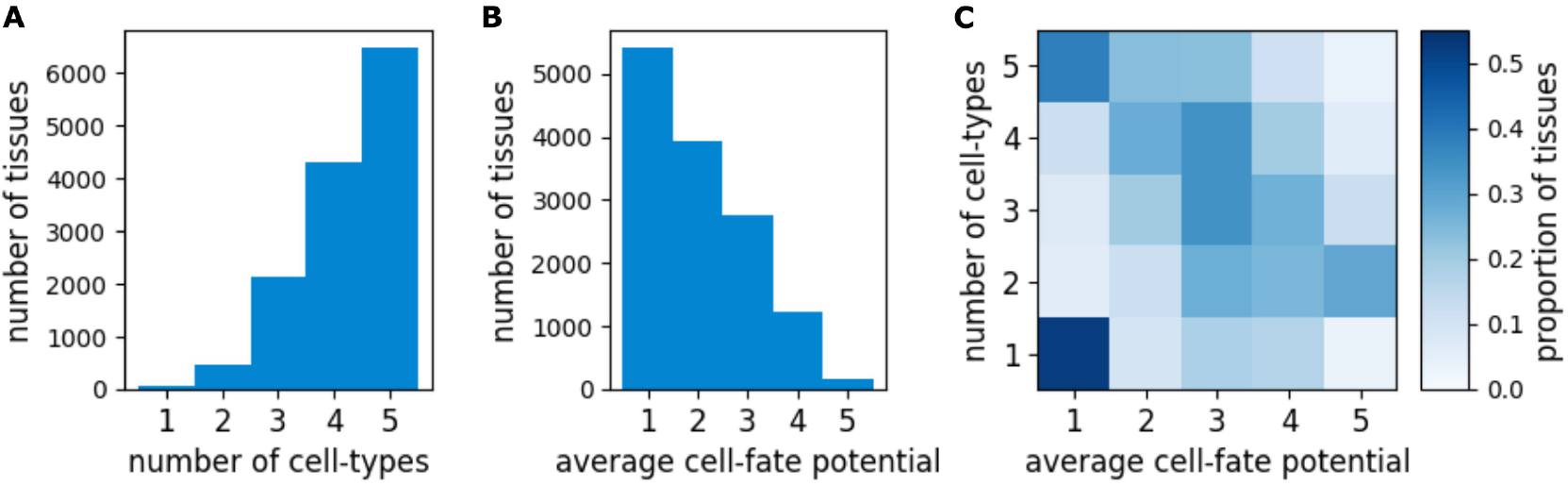
Number of tissue cell-types and their cell-fate potential. **(A)** Histogram of the number of cell-types in model-generated tissues across all parameters. **(B)** Histogram of cell-fate potential averaged across all cell-types in a tissue. **(C)** Heat map of the fraction of tissues with number of cell-types given by the rows and average cell-fate potential given by the columns.

To understand the nature of tissue cell-types, we calculate their cell-fate potentials as the set of all distinct cell-types which it differentiates into across all possible cellular neighborhoods (Fig.2(B), Supplementary Information:FigS2). Note that here cell-fate potential refers to the set of cell-types which any given cell-type *directly* differentiates into. Cell-fate potentials are expected to be wider if one additionally considers indirect differentiation involving multiple transitions. Average cell-fate potential in the model decreases with *P*_*stable*_(*r* = -0.31), and increases with *P*_*diff*_ (*r* = 0.62). Average cell-fate potential also increases with *P*_*den*_(*r* = 0.28): Increasing cellular interaction partners in turn increases the probability that a cell has different fates in different cellular neighborhoods.

Overall, model-generated tissues with more cell-types tend to have lower average cell-fate potential (Fig.2(C), Pearson’s *r* between number of tissue cell-types and average cell-fate potential = -0.35). That is, most model-generated tissues tend to be rich in cell-types and the constituent cell-types tend to have low plasticity. Nevertheless, 31% (57%) of model-generated tissues contain at least one cell-type which can directly (through multiple transitions) differentiate into all 5 tissue cell-types (Supplementary Information:FigS3). These highly plastic cell-types in the model’s tissues are reminiscent of stem cells in regenerative tissues in real organisms (Fuchs and Blau, 2020).

### Tissues vary widely in the fullness and contiguity of cellular arrangements

#### Tissues are organized into domains

In nature, cells of different types are not randomly scattered in a tissue; rather, they are arranged into distinct, but functionally and developmentally interconnected domains. Each tissue domain has a defined cellular composition and sits in a specific position with respect to other tissue domains.

Model-generated tissues resemble natural tissues and are organized into domains. 77% of tissues reach steady states where each cell is part of a stable cellular neighborhood (Supplementary Information: FigS1). Since cellular neighborhoods also encompass cells in adjacent grid-squares, at steady state, contiguous grid-squares tend to be arranged into domains defined by their cellular compositions (Fig.1(E)). We characterize tissues using five features that describe the spatial arrangement of cells and domains (Methods:Analysing final tissues, Supplementary Information:FigS4):

1. *Fullness*, or the number of grid-squares in the final tissue that are occupied by cells, describes tissue size and sparsity (Fig.3(C)).
2. The *number of distinct tissue domains* describes how cells of different types are distributed into distinct stable cellular neighborhoods.
3. *Coverage equality* describes whether or not the tissue is dominated by a domain of one particular cellular composition, as opposed to being more uniformly covered by different domains.
4. *Dispersity* describes whether domains of identical cellular composition are present as a single connected region, or are scattered across the tissue in smaller disjoint regions (Fig.3(B)).
5. *Mixedness* describes the diversity of cellular compositions of grid-squares adjacent to any given grid-square. For example, grid-squares at the edges of domains are more *mixed* than those in the center; in this sense, all disperse tissues are expected to have a high overall mixedness, but some non-disperse tissues with large but intertwined domains (resembling noodles) also have high *mixedness*.

**Figure 3.**
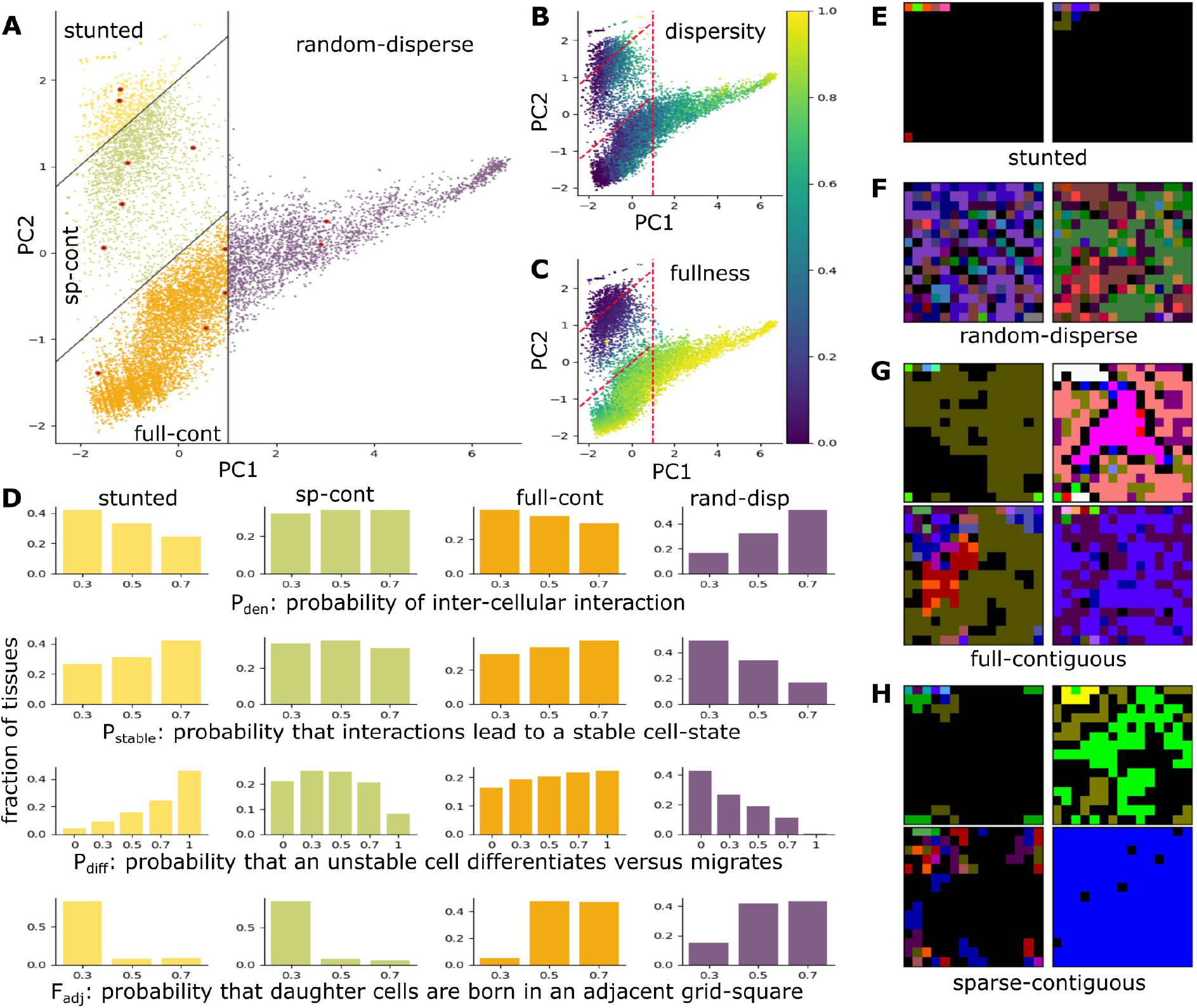
Sectors define qualitatively distinct tissues. **(A)** Spread of model-generated tissues across PC1 and PC2. Each point represents the state of a model-generated tissue at the final time-point. Tissues belonging to the 4 different sectors are indicated by different colors (yellow = stunted, green = sparse-contiguous, orange = full-contiguous, purple = random-disperse). Lines indicate borders between sectors. Red points represent examples of tissues from each sector shown in (E-H). **(B)** Values of *dispersity* across model-generated tissues. **(C)** Tissue *fullness*, which we calculate as the number of occupied grid-squares, across model-generated tissues. **(D)** Histograms for model parameters (rows) encoding tissues belonging to different sectors (columns). **(E-H)** Representative model-generated tissues belonging to different sectors: **(E)***Stunted* tissues show very little growth; **(F)** the *random-disperse* sector contains tissues in which the grid is densely populated by small contiguous domains. **(G)** the *full-contiguous* sector contains tissues in which the grid is densely populated by large contiguous domains; **(H)** the *sparse-contiguous* sector contains diverse forms of tissues. Depending on the distance of the tissue from the sector boundaries, it can resemble tissues in the stunted sector (top-left image), or in the full-contiguous sector (bottom-right image), or be a sparse and disperse tissue (bottom-left image). Importantly, this sector contains tissues where the grid is sparsely populated, yet the domains are contiguous.

Of these five features, *dispersity* and *mixedness* are related to clinically relevant quantities that are calculated from spatial omics studies (Feng et al., 2023).

#### Model-generated tissues fall into four sectors

We perform a Principal Components Analysis of all model-generated tissues against these five features and find that the first two Principal Components (PC1 and PC2) capture 86% of the variance in the data (Methods:Analysing final tissues, Supplementary Information:FigS5, FigS6). Correlations between the PCs and the five tissue features indicate that tissues mostly vary in *dispersity* and *fullness* (Fig.3(B,C)).

We also find that model parameters are significantly predictive of tissue spatial properties: we can divide the plane described by PC1 and PC2 into four sectors such that for 126 out of the 135 parameter sets tested, at least 50 out of 100 tissues independently generated using a single parameter set lie within a single sector (Fig.3(A), black lines). Note that the Principal Components themselves do not separate these tissues into distinct clusters, and tissue properties vary more or less continuously (Supplementary Information:FigS7). Our separation of tissues into sectors is for instructive purposes and for ease of discussion.

Sectors capture qualitative features of tissue morphology and spatial distributions of tissue domains; appropriately, we name these four sectors *stunted* (17% of tissues, Fig.3(E)), *random-disperse* (19% of tissues, Fig.3(F)), *full-contiguous* (48% of tissues, Fig.3(G)) and *sparse-contiguous* (16% of tissues, Fig.3(H)).

Our model parameters are based on local cellular interactions, nonetheless they provide clues about global cellular spatial arrangements: for example, the *stunted* sector is populated by tissues whose cells tend to be non-interactive (low *P*_*den*_) and intrinsically stable (high *P*_*stable*_). Additionally, growth of these tissues is hampered because daughter cells tend to be born within the same grid-square as the mother cell rather than an adjacent square (low *F*_*adj*_) and the levels of cell migration are low (high *P*_*diff*_) (Fig.3(D), first column). In contrast, the *random-disperse* sector is populated by tissues with highly interactive and unstable cells that are highly migratory and tend to produce daughter cells in adjacent grid-squares (Fig.3(D), fourth column). Model parameters that generate *sparse-contiguous* and *full-contiguous* tissues are more moderately distributed than in the *stunted* and *random-disperse* sectors (Fig.3(D), second and third columns).

### Tissue development resembles a sequence of sector crossings

In real organisms, tissue development is expressed as a sequence of intertwined morphogenetic and cell-fate changes (Thowfeequ and Srinivas, 2022; Armenta-Medina et al., 2021). In our model, the four sectors capture tissue morphology as well as cellular compositions and spatial arrangements of tissue domains. Therefore, we project the states of growing tissues onto the two Principal Components described earlier (Fig.3(A)) and express tissue development as sequences of sector crossings (Fig.4, Methods:Analysing tissue developmental trajectories, Supplementary Information:FigS8).

**Figure 4.**
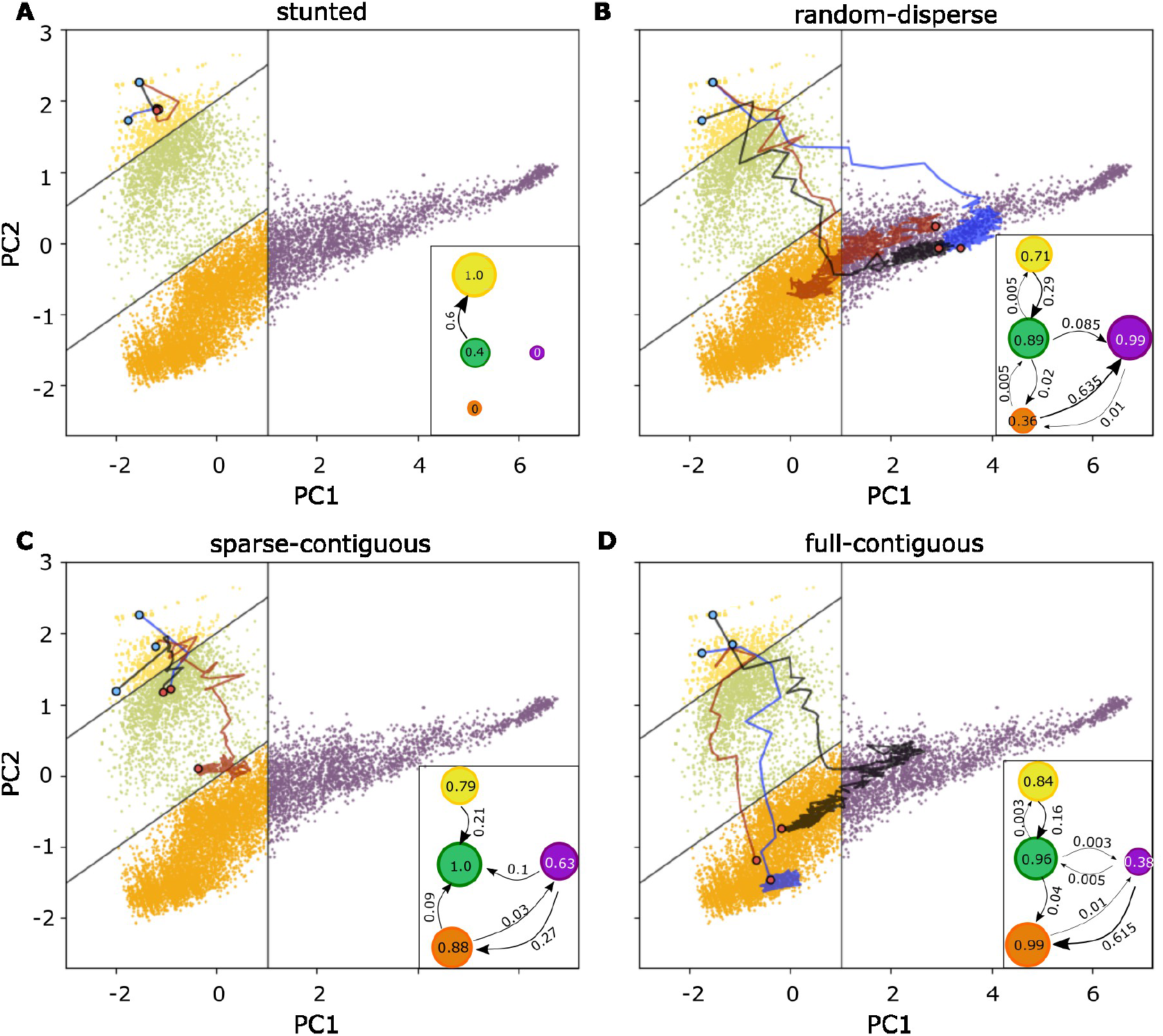
Tissue growth trajectories and sector crossings. Representative tissue growth trajectories projected onto PC1 and PC2 for tissues whose final states fall into (A) *stunted*, (B) *random-disperse*, (C) *sparse-contiguous*, (D) *full-contiguous* sectors. Blue circles represent the initial state of tissues; red circles represent final states of tissues; colored lines indicate growth trajectories. Insets: Graphs indicating probabilities of sector crossings. Nodes (colored circles) represent different sectors (yellow = stunted, green = sparse-contiguous, orange = full-contiguous, purple = random-disperse). Numbers on edges (arrows) indicate the fraction of updates in a trajectory that take a tissue from the source to the target sector averaged across all trajectories for (A) *stunted*, (B) *random-disperse*, (C) *sparse-contiguous*, (D) *full-contiguous* tissues. Numbers within circles represent the average fraction of updates that stay within the corresponding sector.

We summarize tissue developmental trajectories of different tissue types (stunted, full-contiguous, sparse-contiguous and random-disperse) as graphs whose nodes represent sectors and edges represent sector crossings (Fig.4, insets). The edges of the graph are weighted to reflect the fraction of developmental update steps where a particular sector crossing occurs averaged across all tissues of a given tissue-type. For example, we see from the inset in Fig.4(D) that all *full-contiguous* tissues have a *sparse-contiguous* intermediate stage. A majority of these tissue developmental trajectories involve only two sector crossings: *stunted* → *sparse-contiguous* → *full-contiguous*, with multiple update steps within each of the intermediate sectors (e.g. Fig.4(D), red and blue trajectories). A small number of trajectories transition via the *random-disperse* sector and some of these trajectories involve oscillations between the *random-disperse* sector and the final *full-contiguous* sector (e.g. Fig.4(D), black trajectory).

### Full and contiguous tissues heal well

#### Simulating tissue injury and recovery

We test the healing ability of model-generated tissues by simulating injuries to final time-step tissues and assessing their ability to return to their original states (Methods:Analysing tissue healing). We perform three types of injuries:

##### type A

*Single square-all cells* – we ablate all cells of any cell-type from a single randomly chosen grid-square (Fig.5(A), first row).

**Figure 5.**
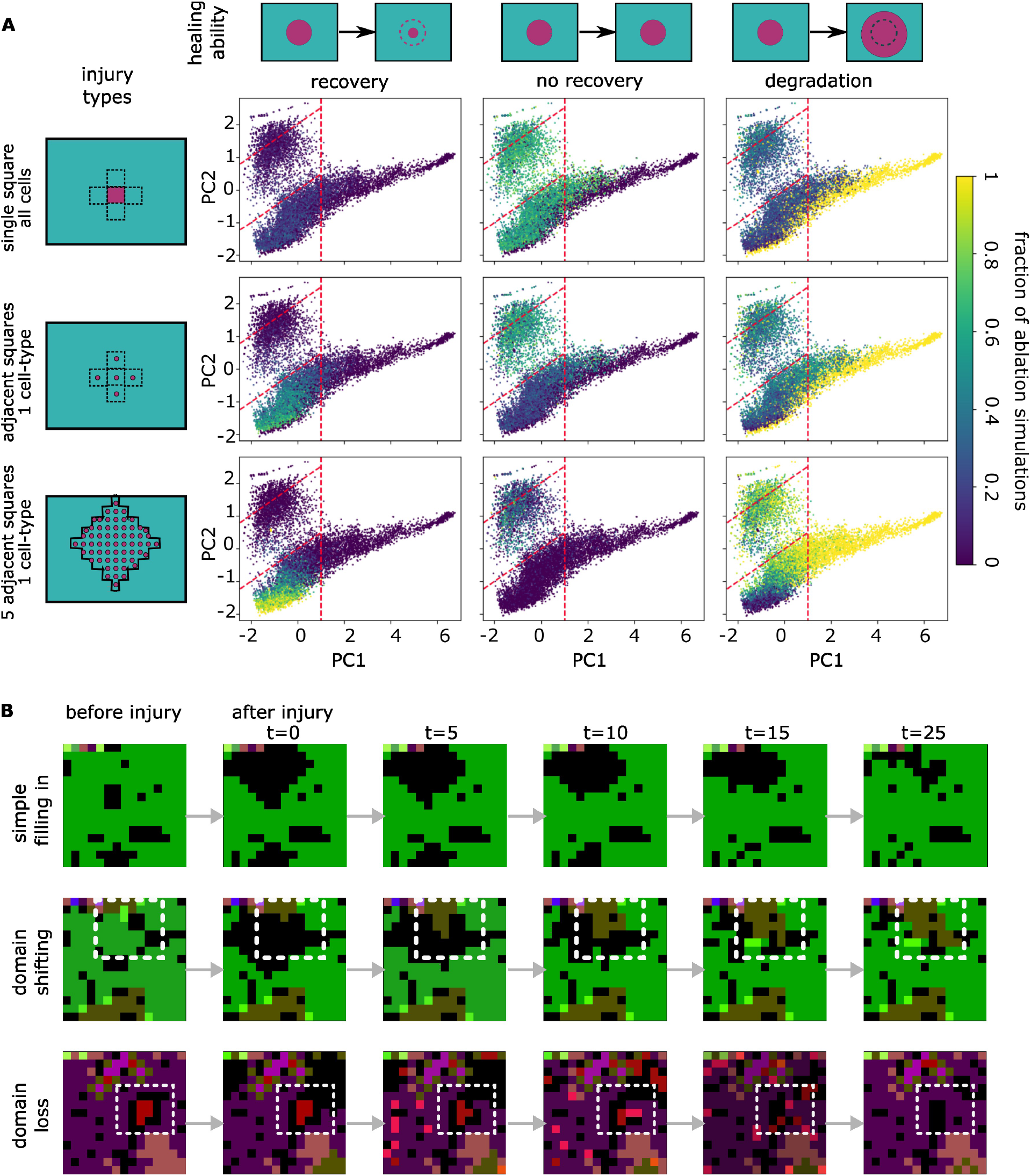
Tissue responses to injury. (A) Tissue healing ability. Points in the scatter plots indicate distinct tissues. Colors of points indicate the fraction of injuries for which the tissue showed recovery (left column), no recovery (middle column) or degradation (right column). Rows indicate injury type: first row – all cells are ablated from a single random grid-square, second row – all cells of a single random cell-type are ablated from a randomly chosen set of adjacent grid-squares, third row – all cells of a single random cell-type are ablated from a randomly chosen set of adjacent grid-squares. (B) Tissue recovery trajectories. We demonstrate three distinct modes of tissue recovery prevalent in the model for type C injuries: first row – the ablated cell-type is replenished through cell division in adjacent grid-squares, second row – the injury occurs at a boundary between two domains (white box), thus promoting cell division in both domains, ultimately leading to a shifted domain boundary in the tissue after healing, third row – the injury affects cell fates throughout the tissue, ultimately destabilizing a domain (red domain in white box) leading to its loss.

##### type B

*Adjacent squares-single cell-type* – we ablate all cells of a single randomly chosen cell-type from all grid-squares adjacent to a single randomly chosen grid-square. This amounts to a single cell-type being ablated from 5 contiguous grid-squares (Fig.5(A), second row).

##### type C

*5 neighbors-single cell-type* – we ablate all cells of a single randomly chosen cell-type from all grid-squares that can be reached in ≤ 5 steps from a randomly chosen grid-square. This amounts to a single cell-type being ablated from 61 contiguous grid-squares, which is 27% of the tissue’s body (Fig.5(A), third row).

After injuring a tissue, we allow it to heal by updating the injured tissue for 100 time-steps using the same Cell Fate Rules originally used during tissue development (see Methods:Analysing tissue healing). We assess the ability of a tissue to heal by comparing the Euclidean distance between the state of the tissue before injury and immediately after injury (*IM*_*t*=0_) against the distance between the state of the tissue before injury and after 100 time-steps of healing (*IM*_*t*=100_). We say that a tissue heals if *IM*_*t*=100_*< IM*_*t*=0_; degrades if *IM*_*t*=100_*> IM*_*t*=0_; does not recover if *IM*_*t*=100_ = *IM*_*t*=0_. Notably, cell migration and cell division are probabilistic decisions in our model, therefore tissues are not expected to heal perfectly and do not achieve the exact configuration before injury (Fig.5(B)). We perform 25 independent injuries of type A and type B for each tissue and 10 independent injuries of type C for each tissue and report the fraction of injuries for which the tissue heals (*recovery-fraction*: Fig.5(A), first column), does not recover (*no-recovery-fraction*: Fig.5(A), second column), or degrades (*degradation-fraction*: Fig.5(A), third column).

In order to gain some intuition for the process of tissue healing, consider how cell-fate in the model depends on the cellular neighborhood: For cells present in the middle of a domain, ablation of one or more cells in a single adjacent grid-square does not change its neighborhood, and thus does not influence its behavior. Therefore, injuries affecting cells in a single grid-square, such as type A injuries, are typically not sufficient to evoke a healing response; these injuries only perturb the tissue if the affected grid-square is at the edge of a domain. In keeping with this intuition, the *degradation-fraction* for type A injuries is highest in tissues with high *mixedness*, in which a majority of grid-squares lie on the edges of domains (Pearson’s *r* between *degradation-fraction* and *mixedness* for type A injuries = 0.51 (p-value = 0.0)). Overall, for type A injuries, 61.5% of all tissues tend to neither heal nor get worse, and the *no-recovery-fraction* is greater than 0.5 (green dots in Fig.5(A) second column, row 1).

In contrast, type B and type C ablations are bound to change cellular neighborhoods for cells occupying grid-squares in the middle of the ablation. We find that the ability to heal from type B and type C injuries was enriched in the *full-contiguous* sector: For type B injuries, only 16.8% of all tissues have a recovery-fraction ≥ 0.5, and 99.4% of these tissues are in the *full-contiguous* sector (Supplementary Information:FigS9). For type C injuries, 24.7% of all tissues have a recovery-fraction ≥ 0.5, and 97.1% of these tissues are in the *full-contiguous* sector. These results reveal a strong connection between tissue morphology and healing, and highlight evolutionary constraints on tissue architecture stemming from the requirement of tissues to heal. We expect that these constraints apply universally to all multicellular groups despite their independent origins (Birnbaum and Alvarado, 2008).

### Mechanism of tissue recovery in the model

In order to understand the general mechanism of tissue healing in the model, we tested how model parameters are correlated with *recovery-fractions*: for both type B and type C injuries, the ability to heal shows the strongest correlation with *F*_*adj*_(*r* = 0.44 for type B and *r* = 0.30 for type C, see Supplementary Information:FigS10). This suggests that the mechanism of tissue healing in the model depends on cell division in adjacent grid-squares that replenishes lost cells in the injury sites (for example Fig.5(B),rows 1,2). This mechanism is notably similar to that seen in mammalian tissues (Clevers, 2013; Wei et al., 2021), as well as during tissue healing in plants, where cells adjacent to injuries switch their cell division axes in order to replace lost cells (Marhava et al., 2019).

## DISCUSSION

We present here an agent-based spatial model of cell-fate decisions in tissues. Notably, our model trades the ability to explicitly encode the exact cellular neighbors and cell shapes that are captured in more traditional models (Farhadifar et al., 2007; Marée et al., 2007; Bock et al., 2010) for simplicity, generality, and the ability to survey vast spaces of tissue developmental rules. We use our model to explore the diversity of tissue architectures produced by simple developmental rules and the association between tissue architecture and tissue healing. We categorize model-generated tissues into four sectors – *stunted, sparse-contiguous, full-contiguous* and *random-disperse* – which represent tissue morphology as well as cellular compositions. These sectors capture a wide variety of tissue architectures: For example, most natural tissues tend to be contiguous and ordered, such as animal epithelia (Buckley and St Johnston, 2022), mammalian striated muscle bundles (Mukund and Subramaniam, 2020), and plant leaf epidermis (Zuch et al., 2022), which are examples of tissues resembling *full-contiguous* tissues. In contrast, the leaf spongy parenchyma forms a sparse but contiguous tissue (Borsuk et al., 2022) and resembles tissues in the *sparse-contiguous* sector, while animal blood resembles model-generated *random-disperse* tissues (Nagahata et al., 2022).

We characterize tissue growth in the model in terms of crossings between different sectors, which recapitulates the manner in which tissue development in real organisms is described, where morphological changes are coupled with cell-fate changes. For example, during human embryo development, compaction of the ball of cells during Morula is coupled with cellular differentiation into the inner cell mass and the trophectoderm, and subsequent cavitation of the ball of cells is coupled with differentiation of the inner cell mass into the epiblast and the hypoblast (Thowfeequ and Srinivas, 2022). These cell-fate decisions are not cell autonomous; instead, all through this process, different cell-types communicate with each other and play an important role in regulating each other’s cellular decisions (Okubo et al., 2023).

Importantly, we find a strong association between model-generated tissue structures with their ability to regenerate; particularly, the ability to heal is enriched in *full-contiguous* tissues. Thus our results indicate that tissue healing could act as a selective pressure that moulds tissue architecture. Moreover, the mechanism of healing in the model is through replacement of damaged cells by cell-division in adjacent regions; this resembles the process of healing in plant (Marhava et al., 2019) as well as animal tissues (Clevers, 2013; Wei et al., 2021). However, our model is statistical in nature; therefore, it identifies the most probable mechanisms across a wide range of modes of cellular interaction. Thus outputs of our model may not match specific mechanisms in particular organisms: For example, regeneration in planarian flatworms depends on widely circulating neoblast cells (Reddien, 2018), which is in contrast to the local mechanism enriched in the model.

Overall, our spatial model of cell-fate captures essential features of tissue growth, form and function. Our model also maps how varying local intercellular interactions can lead to changes in global tissue architecture, which in turn is closely related to the ability of tissues to heal. These inter-related tissue properties allow experimental tests of the model’s predictions. We highlight here one such experimental test: our model predicts that increasing the density of inter-cellular interactions should lead to more disperse tissues (Fig.3(D), first row), and disperse tissues tend to have low healing capacity (Fig.5). To experimentally test this prediction, one could tune the density of cellular interaction by manipulating the types of receptors expressed on tissue cell-types and observe the ensuing effects on tissue architecture and tissue healing. Additionally, our framework is flexible and can be modified to encode special features of particular tissues; we highlight two natural extensions: First, while two- and three-dimensional grids may be appropriate to study many solid tissues, tissues such as the plant spongy mesophyll (Borsuk et al., 2022), or the branched animal lung epithelium (Schittny, 2017) can be more faithfully represented with more irregular geometries. Second, the model can be expanded by including external signals associated with different grid locations irrespective of their cellular compositions, which could represent morphogen gradients that form an integral aspect of animal (Romanova-Michaelides et al., 2022) as well as plant development (Hu et al., 2021).

## METHODS

### Sampling the space of tissue developmental parameters

We simulate tissues with *N* = 5 cell-types in a *G* × *G* 2-dimensional grid with periodic boundaries, where *G* = 15. The geometry of the tissue is described by the grid’s adjacency matrix A, where *A*(*g*_*i*_, *g*_*j*_) = 1 if grid-squares *g*_*i*_ and *g*_*j*_ are adjacent to each other. In a 2-dimensional grid, each square is adjacent to 4 other squares, as shown in Fig.1(B). We uniformly survey parameter values from the following sets: *P*_*den*_ ∈ {0.3, 0.5, 0.7}, *P*_*stable*_ ∈ {0.3, 0.5, 0.7}, *P*_*diff*_ ∈ {0, 0.3, 0.5, 0.7, 1} and *F*_*adj*_ ∈ {0.3, 0.5, 0.7}. For each combination of parameter values, we draw 100 replicate *tissue development rules*, and overall, we examine tissues resulting from 13,500 distinct rules. For 77% of these simulated tissues, cellular compositions of all grid-squares reach steady state within 400 time-steps.

### Generating tissue developmental rules

#### Interaction matrix

The interaction matrix *I* is a matrix of size *N* × *N* where

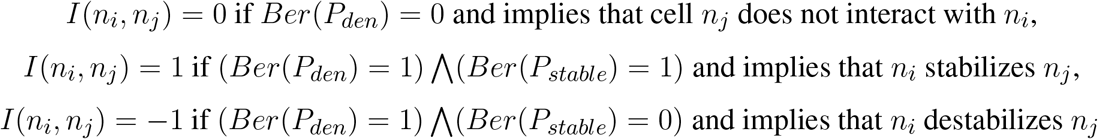

Where *Ber*() symbolizes a Bernoulli random variable. Note that the interaction matrix for each tissue is randomly generated according to parameters *P*_*den*_ and *P*_*stable*_ and remains fixed for the duration of the simulation.

#### Cell Fate Rules

Let *H*^*a*^ = 0, 1^*N*^ be an N-length binary vector describing an arbitrary cellular neighborhood, such that *H*^*a*^(*n*) = 1(0) indicates the presence (absence) of cell-type *n* in neighborhood *H*^*a*^. An N-length vector *Stable*^*a*^ encodes the stability of cell-types present in the neighborhood *H*^*a*^.

*Stable*^*a*^(*n*_*i*_) = 1 and cells of cell-type *n*_*i*_ are stable in neighborhood *H*^*a*^ if:

- it is intrinsically stable and receives no destabilizing interactions: that is, *I*(*n*_*i*_, *n*_*i*_) = 0*or*1, and ∄*n*_*j*_*s*.*t*.(*H*^*a*^(*n*_*j*_) = 1) ⋀ (*I*(*n*_*j*_, *n*_*i*_) = −1)
- it is intrinsically unstable and receives stabilizing interactions: that is, *I*(*n*_*i*_, *n*_*i*_) = −1, and ∃*n*_*j*_*s*.*t*.(*H* ^*a*^(*n*_*j*_) = 1) ⋀ (*I*(*n*_*j*_, *n*_*i*_) = 1)

*Stable*^*a*^(*n*_*i*_) = −1 and cells of cell-type *n*_*i*_ are unstable in neighborhood *H*^*a*^ if:

- it is intrinsically stable and receives destabilizing interactions: that is, *I*(*n*_*i*_, *n*_*i*_) = 0*or*1, and ∃*n*_*j*_*s*.*t*.(*H*^*a*^(*n*_*j*_) = 1) ⋀ (*I*(*n*_*j*_, *n*_*i*_) = −1)
- it is intrinsically unstable and receives no stabilizing interactions: that is, *I*(*n*_*i*_, *n*_*i*_) = −1, and ∄*n*_*j*_*s*.*t*.(*H*^*a*^(*n*_*j*_) = 1) ⋀ (*I*(*n*_*j*_, *n*_*i*_) = 1)

*Stable*^*a*^(*n*_*i*_) = 0 for cell-types not present in neighborhood *H*^*a*^. Unstable cells either differentiate or migrate. For cells present in *H*^*a*^, we encode Cell Fate Rules as a pair of look-up tables: A differentiation table *Diff* ^*a*^ and a migration table *Mig*^*a*^.

- *Diff* ^*a*^(*n*_*i*_) = *n*_*j*_ indicates that cell-type *n*_*i*_ differentiates into cell-type *n*_*j*_ in neighborhood *H*^*a*^. If (*Stable* (*n*_*i*_) = −1) ⋀ (*Ber*(*P*_*diff*_) = 1), *n*_*j*_ is randomly drawn from among the N cell-types. On the other hand, if (*Ber*(*P*_*diff*_) = 1) ⋁ (*Stable*^*a*^(*n*_*i*_) = 1), then *n*_*j*_ = *n*_*i*_.
- *Mig*^*a*^(*n*_*i*_) = *m* such that *m* ≤ *G* indicates that cell-type *n*_*i*_ in neighborhood *H*^*a*^ migrates to a grid-square a distance m away from its current grid-square. If (*Stable*^*a*^(*n*_*i*_) = −1) ⋀ (*Be r*(*P*_*diff*_) = 0), *m* is uniform randomly drawn from 0, 1, 2, …*G*. On the other hand, if (*Ber*(*P*_*diff*_) = 1) ⋁ (*Stable*^*a*^(*n*_*i*_) = 1), then *m* = 0.

Note that although Cell Fate Rules for cellular differentiation and cell migration are randomly generated according to the parameter *P*_*diff*_, they remain fixed for the duration of the simulation.

### Updating tissue states

At any time-step, the state of the tissue is completely characterized by the cellular compositions of its grid-squares: 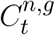 gives the number of cells of cell-type *n* for *n* ∈ 1, 2, 3, …*N* in grid-square *g* for *g* ∈ 1, 2, 3, …*G*^2^ at time-step *t*. At *t* = 0, we initialize the tissue by randomly placing 20 cells of random cell-types across the connected set of grid-squares 0, 1, 2, 3, 4. At each time-step of the simulation, the state of the tissue is updated through sequential steps of cell death, cell division, and cell differentiation and migration.

#### Cell death

In the tissue at time *t*, the total number of cells in a grid-square *g* is given by 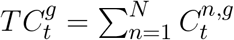. Then the probability of cell-death in this grid-square is given by

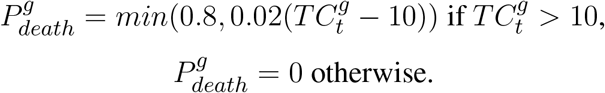

That is, there is no cell-death in grid-squares with 10 or fewer cells, and the probability that any cell dies increases linearly as the number of cells in the grid-square increases beyond 10.

We draw the number of cells of cell-type *n* in grid-square *g* that die in this time-step, 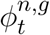 from the binomial distribution 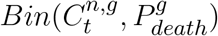. Then, 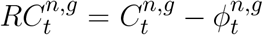 gives the cells of cell-type *n* that remain in grid-square *g*.

#### Cell division

After cell death, the total number of cells remaining in some grid-square *g*_*i*_ is given by 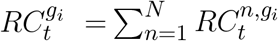. Then the probability of cell-division in this grid-square is given by 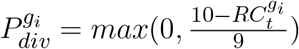. That is, there is no cell-division in unoccupied grid-squares and in squares with 10 or more cells, and the probability that any cell divides decreases linearly as the number of cells in the grid-square increases up to 10.

We draw the number of cells of cell-type *n* in grid-square *g*_*i*_ that divide in this time-step, 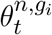 from the binomial distribution 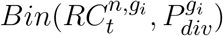. 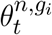 is then the number of daughter cells of cell-type *n* produced in grid-square *g*_*i*_.

A fraction (1 − *P*_*adj*_) of these daughter cells are born within grid-square *g*_*i*_. The rest of the daughter cells are assigned to randomly chosen grid-squares adjacent to *g*_*i*_. Let 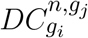 be the number of daughter cells of cell-type *n* produced by division of cells in grid-square *g*_*i*_ that are born in grid-square *g*_*j*_. For any *g*_*j*_ not adjacent to *g*_*i*_, 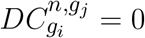.

Then, 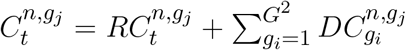 gives the total number of cells of type *n* in grid-square *g*_*j*_ after cell-division.

Note that in the model, the probability of cell-division is the same for all cells of all cell-types in a given grid-square. Additionally, two cells of the same cell-type in the same grid-square may divide differently: Depending on 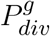, it is possible that only one of these cells divides. In case both these cells divide, depending on 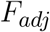, it is also possible that the daughters are produced in different grid-squares.

#### Cell differentiation and migration

Let N-length binary vector 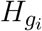 represent the cellular neighborhood for grid-square *g*_*i*_ in *C*_*t*_.

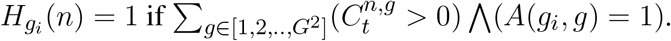

Cell migration and differentiation rules for cells in grid-square *g*_*i*_ can then be calculated using the Cell Fate Rule 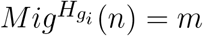: cells of type *n* migrate from grid-square *g*_*i*_ if *m >* 0, else not.

In order to find the new locations of cells migrating from grid-square *g*_*i*_, we use matrix powers of the adjacency matrix, *A*^*m*^: The set of all grid-squares m steps away from *g*_*i*_ is 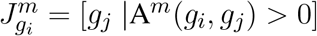. For each cell in *g*_*i*_ that migrates a distance m away, we randomly choose its new grid-square from 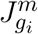. We encode updated locations of migrating cells in *µ*: number of cells of cell-type *n* originally in grid-square *g*_*i*_ which have migrated to grid-square *g*_*j*_ is given by 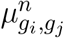.

Note that *µ* only contains the new locations of cells that migrate, i.e., for all cells of any cell-type *n* such that 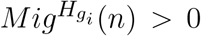. Non-migrating cells are either stable in the current cellular neighborhood 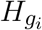, or they differentiate. Cell Fate Rules governing the fate of non-migrating cell-types are given by 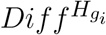. We encode updated cell-types for non-migrating cells in 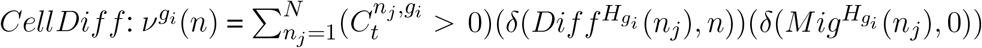. That is, 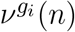 is the number of non-migrating cells of any cell-type in *g*_*i*_ which differentiate into cell-type *n*.

The state of the tissue in the next time-step is then given by:

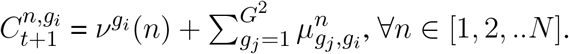

We synchronously update each grid-square at each model time-step, and perform 500 updates for each tissue.

### Analysing final tissues

We characterize tissue morphology based on how tissue domains (regions of uniform cellular composition within tissues) are arranged relative to each other. In order to do this, we measure five quantities:

#### Fullness

We calculate tissue fullness as the number of grid-squares that are occupied by cells. This is then given as 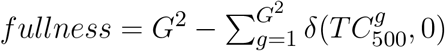.

#### Number of unique domains

We express the cellular composition of any occupied grid-square as a N-length binary vector *κ*^*g*^, where *κ*^*g*^(*n*) = 1 if 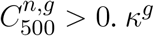 can take on one of 2^*N*^ values depending on the presence or absence of the N cell-types. We arrange these 2^*N*^ distinct cellular compositions according to the standard binary ordering. Tissues in the model tend to be composed of contiguous domains of uniform cellular composition, such that the cellular compositions of grid-squares within a single domain are identical to each other. Therefore, the number of unique cellular compositions (*UD*) across grid-squares gives us the number unique tissue domains (*NUD*). In this work, the number of cell-types *N* = 5, therefore the maximum value of *NUD* = 2^5^, and *NUD* = 1 for completely uniform tissues with a single domain.

#### Coverage equality

Let *DS* be an *NUD* length vector where *DS*(*i*) reflects the size of the *i*^*th*^ domain, and is given by the number of grid-squares with cellular composition identical to the *i*^*th*^ unique domain. We say that a tissue has high *coverage equality* if each unique domain cellular composition is represented by roughly equal number of grid-squares; i.e., 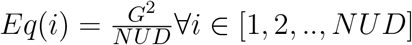. At the other end, for a tissue with extremely unequal distribution of cellular compositions across the grid, *Uneq*(1) = *G*^2^ and *Uneq*(*i*) = 0*∀iin*[2, 3, .., *nud*]. We calculate *coverage equality* of the tissue as:

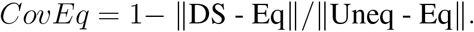

#### Dispersity

Any set of grid-squares of identical composition, say *UD*(*i*), is either completely contiguous, or splits into smaller subsets of contiguous regions. Contiguity of grid-squares can be assessed using the adjacency matrix *A*: A subset of grid-squares is contiguous if it forms a connected component, i.e., any grid-square in the subset can be reached from any other grid-square in the subset by traversing the grid. We encode the number of contiguous components with cellular compositions identical to *UD*(*i*) as *DComp*(*i*). We call a tissue *disperse* if grid-squares with identical cellular compositions are split into multiple distinct connected components. We measure the dispersity of domains with a cellular composition *UD*(*i*) as 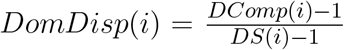. We measure *dispersity* of the tissue as the average *DomDisp*; i.e., 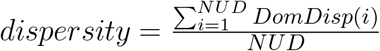.

#### Mixness

Finally, we examine the local neighborhood of each grid-square by counting *GridMix*_*i*_: the number of unique cellular compositions across the set of grid-squares adjacent to *g*_*i*_, including itself. For a 2-dimensional grid, *GridMix*_*i*_ is at most 5. We describe a *G*^2^ length vector *SqMix*, where 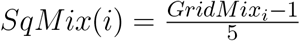, such that for a grid-square in the middle of a contiguous domain, *SqMix*(*i*) = 0. We also assign *SqMix*(*i*) = 0 if grid-square *g*_*i*_ is unoccupied. We define tissue *mixedness* by averaging *SqMix* over all grid-squares.

We characterize the morphology of any model-generated tissue in the final time-point by the set of five numbers: *fullness, NUD, CovEq, dispersity* and *mixness*.

#### Principal Components and sectors

These 5 properties are not independent of each other; particularly, *mixness* and *dispersity* are closely related quantities (Pearson’s correlation coefficient = 0.74). To find relevant dimensions across which properties of model-generated tissues in the final time-point (*t* = 500) vary, we perform Principal Components Analysis. We find that the first two Principal Components (PC1 and PC2) explain 85.6% of the variance, and we now characterize tissues based on PC1 and PC2.

In order to assess how tissue properties (PC1 and PC2) change with model parameters, and for ease of discussion, we further split the space of tissues into *sectors*, such that tissues generated using the same set of model parameters (*P*_*den*_, *P*_*stable*_, *P*_*diff*_, *F*_*adj*_) are likely to all fall within the same *sector*. Based on our raw assessment of model-generated tissues, we choose to divide the space of tissues into *four* sectors. Sector boundaries are straight lines: boundaries between the *stunted* and *sparse-contiguous*, and the *sparse-contiguous* and *full-contiguous* sectors are characterized by their slopes and y-intercepts. The boundary separating the *random-disperse* sector is vertical, and is characterized by its x-intercept. We find optimal values for the slopes and intercepts of sector boundaries which maximize the number of parameter sets for which at least 50% of generated tissues lie within the same sector. For the sector boundaries we use, 50% of generated tissues lie within the same sector for 93.3% of the 135 parameter sets sampled in this study.

### Analysing tissue developmental trajectories

For each tissue, for each intermediate time-point *t* during its development, we calculate the five quantities that describe its morphology: *fullness*_*t*_, *NUD*_*t*_, *CovEq*_*t*_, *dispersity*_*t*_ and *mixness*_*t*_. We scale each of these properties: For example, 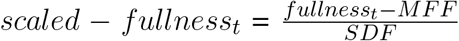; where *MFF* is the average *fullness* across all tissues at *t* = 500, and *SDF* is the standard deviation of *fullness* across all tissues at *t* = 500. We then project scaled values of properties of tissues at intermediate time-points onto the Principal Components (PC1 and PC2) that describe model-generated tissues in the final time-point. This allows us to place developing tissues at intermediate time-points into the four *sectors*: *stunted, sparse-contiguous, full-contiguous and random-disperse*.

For each tissue *T*_*i*_ such that *i* ∈ [1, 2, …, 13500], let the the sector assignment at time *t* be 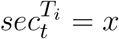, for *x* ∈ [*stunted, sparse-contiguous, full-contiguous, random-disperse*]. We describe tissue developmental trajectories in terms of sequences of sector crossings: 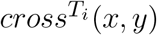 is the number of update steps during its development in which the tissue transitions from a sector *x* to a sector *y*, where *x, y* ∈ [*stunted, sparse-contiguous, full-contiguous, random-disperse*]. For example, 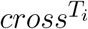 (*stunted, sp* − *contig*) is the number of times *T*_*i*_ transitions from the *stunted* to the *sparse-contiguous* sector, and 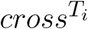 (*sp* − *contig, sp* − *contig*)is the number of update steps in which *T*_*i*_ stays within the *sparse-contiguous* sector.

We then estimate *sector transition probabilities* (*STP*) for each sector: For all tissues which end up in sector *z* ∈ [*stunted, sparse-contiguous, full-contiguous, random-disperse*] at *t* = 500,

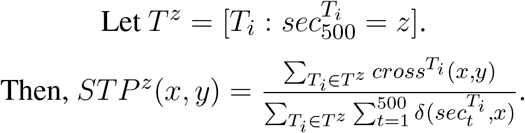

### Analysing tissue healing

#### Producing injuries

For each tissue, we test its ability to heal by simulating three kinds of injuries:

##### type A

For each tissue, we perform 25 independent injuries of this type. For some tissue *T*_*i*_ among the 13500 model-generated tissues, we set the initial state of the injured tissue as the final state of tissue *T*_*i*_, that is:

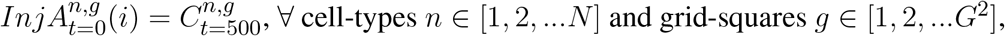

where *i* ∈ [1, 2, …25] represents the injury-id.

Then we ablate all cells of all cell-types from a single grid-square. That is, for some randomly chosen grid-square *g*_*r*_ such that 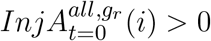, we set 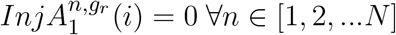.

##### type B

For each tissue, we perform 25 independent injuries of this type. For some tissue *T*_*i*_ among the 13500 model-generated tissues, we set the initial state of the injured tissue as the final state of tissue *T*_*i*_, that is:

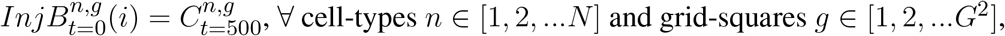

where *i* ∈ [1, 2, …25] represents the injury-id.

In this type of injury, we ablate all cells of one particular cell-type from all grid-squares adjacent to a given grid-square. To do this, for some randomly chosen grid-square *g*_*r*_ and a cell-type *n*_*r*_ such that 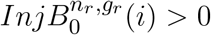, we set 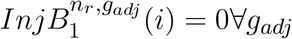 such that *A*(*g*_*r*_, *g*_*adj*_) *<*= 1.

##### type C

For each tissue, we perform 10 independent injuries of this type. For some tissue *T*_*i*_ among the 13500 model-generated tissues, we set the initial state of the injured tissue as the final state of tissue *T*_*i*_, that is:

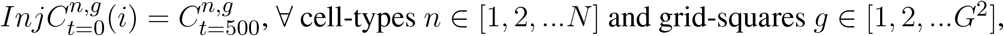

where *i* ∈ [1, 2, …25] represents the injury-id.

In this type of injury, we ablate all cells of one particular cell-type from all grid-squares ≤ 5 steps away from a given grid-square. To do this, for some randomly chosen grid-square *g*_*r*_ and a cell-type *n*_*r*_ such that 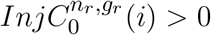, we set 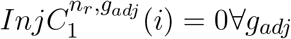 such that *A*(*g*_*r*_, *g*_*adj*_) *<*= 5.

#### Injury Magnitude

Finally, we calculate the magnitude of an injury as the Euclidean distance between the uninjured tissue and the tissue immediately after injury: *IMX*_*t*=1_(*i*) = ∥(*InjX*_*t*=0_(*i*) − *InjX*_*t*=1_(*i*))∥, for injury-ids *i* ∈ [1, 2, …, 25].

#### Tissue recovery

In order to simulate tissue recovery, we update the state of injured tissues for 100 time-steps using the same Cell Fate Rules that were used to simulate tissue development as described above in Updating tissue states. For each tissue, we test how the magnitude of the injury changes during recovery for the 25 replicates of type A and type B injuries, and for the 10 replicates of type C injuries): For the *i*^*th*^ replicate injury, the magnitude of the injury after 100 time-steps of recovery is given by

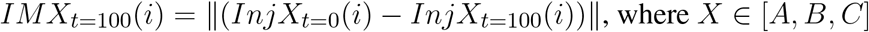

We say that a tissue is from injury i of type X:

- is healing if *IMX*_*t*=100_(*i*) *< IMX*_*t*=1_(*i*).
- shows no recovery if *IMX*_*t*=100_(*i*) = *IMX*_*t*=1_(*i*).
- is degrading if *IMX*_*t*=100_(*i*) *> IMX*_*t*=1_(*i*).

For each tissue, for each injury type, we measure the fraction of replicate injuries out of 25 (10 for type C) that result in healing (*RecFrac*), no recovery (*NoRecFrac*), or tissue degradation (*DegFrac*).

## ACKNOWLEDGEMENTS

This work was supported by the taxpayers of South Korea through the Institute for Basic Science, Project Code IBS-R020-D1. We thank all members of the Living Matter Theory group at IBS-CSLM for engaging discussions during the conception of this work. We thank John McBride for constructive comments during the writing of this manuscript.

## SUPPLEMENTARY INFORMATION

The supplementary file contains the following figures:

- FigS1: Histogram for the number of time-steps to tissue steady-state.
- FigS2: Scatter-plots showing the dependence on model parameters of number of distinct tissue cell-types, average cell-fate potential.
- FigS3: Scatter-plots showing the dependence on model parameters of maximum cell-fate potential of tissue cell-types.
- FigS4: Heatmaps displaying dependence on model parameters of the 5 spatial properties using which we characterize model-generated tissues.
- FigS5: Elbow plot showing the variance in the data explained by each PC
- FigS6: Scatter-plots showing the dependence on model parameters of Principal Components which represent tissue spatial properties.
- FigS7: Hierarchical clustering of model-generated tissues.
- FigS8: Histograms for the number of sector crossings, number of time-steps to reach the final sector.
- FigS9: Scatterplots displaying the extent of tissue healing for tissues in different sectors in response to type B injuries.
- FigS10: Heatmaps displaying dependence of tissue healing in response to type B injuries on model parameters.

## Notes

### Competing Interest Statement

The authors have declared no competing interest.

### Summary of Updates

Updated Abstract. New results in section titled 'Full and contiguous tissues heal well'. New panel added to Fig5.

