## SupplementalSections for "Linking Tissue Morphology and Tissue Healing in a Cell-Fate Model"

March 8, 2026

#### 1 Tissue steady states

Overall, 77% of model-generated tissues reach steady state, where the set of cell-types occupying any grid square does not change upon update, within 400 time-steps (FigS.1). The number of time-steps to steady-state increases with model parameters $P_{den}$ and $F_{adj}$ (Pearson's correlation coefficients: 0.17, 0.21, respectively), and decreases with parameters $P_{stable}$ and $P_{diff}$ (Pearson's correlation coefficients: -0.18, -0.34, respectively).

Across the 4 sectors the average number of time-steps taken by tissues to reach steady state are – *stunted* tissues: 36.9 time-steps, *sparse-contiguous* tissues: 51.3 time-steps, *full-contiguous* tissues: 164.2 time-steps, *random-disperse* tissues: 354.5 time-steps.

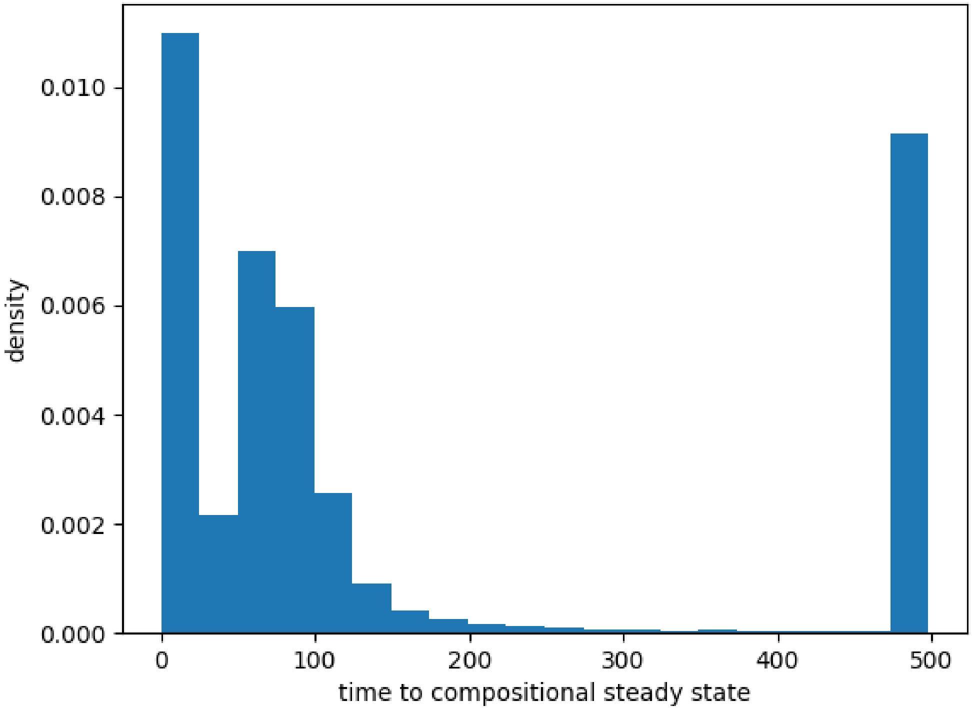


FigS. 1: Histogram for the number of time-steps to tissue steady-state: x-axis represents model time-steps and the y-axis represents the density of model-generated tissues.

#### 2 Tissue cell-types and their cell-fate potentials

Model-generated tissues tend to be rich in cell-types: across tissues, the average number of cell-types is 4.2, and 48% of tissues contain all 5 cell-types. The number of cell-types in tissues increases with parameter $P_{stable}$ (Pearson's correlation coefficient: 0.15), and decreases with parameters $P_{den}$ and $P_{diff}$ (Pearson's correlation coefficients: -0.14, -0.41, respectively)(FigS.2(Row1)). Across the 4 sectors, the average number of cell-types in *stunted* tissues is 4.06, in *sparse-contiguous* tissues is 4.18, in *full-contiguous* tissues is 4.15 and in *random-disperse* tissues is 4.64.

At the same time, average cell-fate potential of tissue cell-types tends to be low: the average number of cell-types any given tissue cell-type directly differentiates into is 2.3. Average cell-fate potential equals 5 for only 1.2% of model-generated tissues. Average cell-fate potential in tissues decreases with parameter $P_{stable}$ (Pearson's correlation coefficient: -0.31), and increases with parameters $P_{den}$ and $P_{diff}$ (Pearson's correlation coefficients: 0.28, 0.62, respectively)(FigS.2(Row-2)). Across the 4 sectors, the average cell-fate potential in *stunted* tissues is 2.51, in *sparse-contiguous* tissues is 2.32, in *full-contiguous* tissues is 2.3 and in *random-disperse* tissues is 2.12.

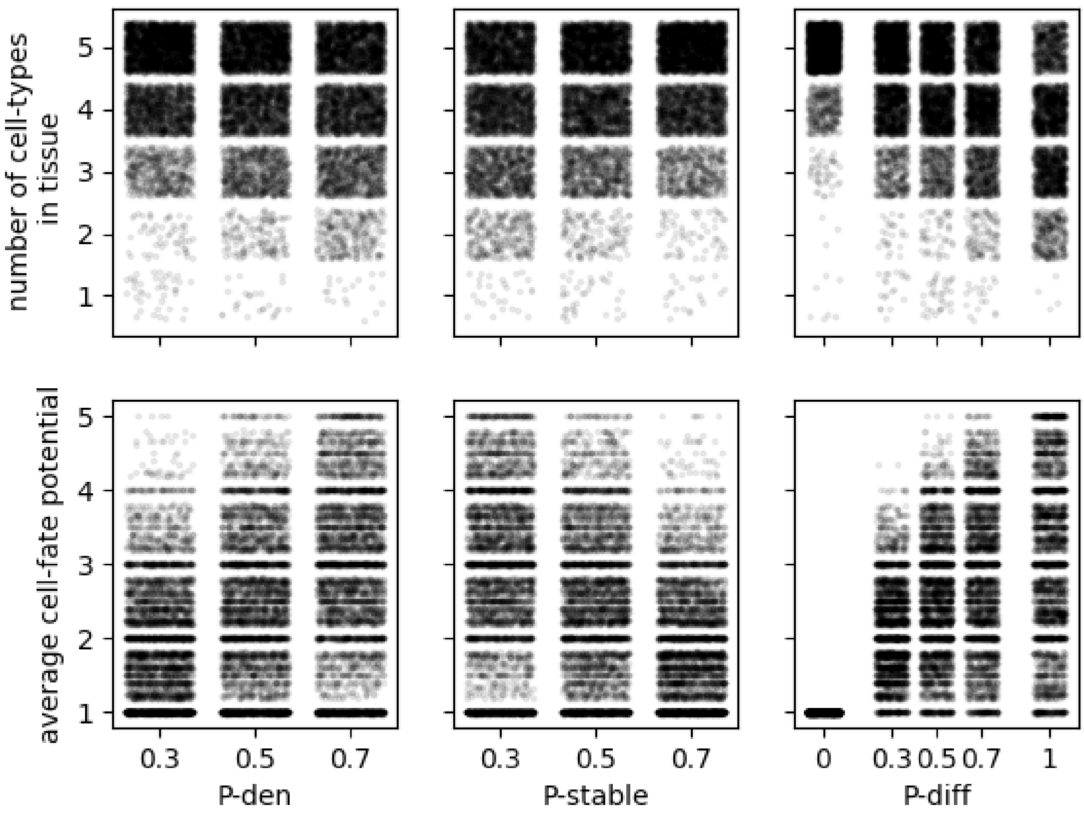


FigS. 2: Scatter-plots showing the dependence on model parameters of (Row-1) number of distinct tissue cell-types, (Row-2) average cell-fate potential. From left to right, the x-axis represents $P_{den}$, $P_{stable}$, $P_{diff}$. Each point represents a model-generated tissue. Noise has been added to the position of points to make density more apparent.

Across tissues, maximum cell-fate potential (due to direct differentiation) for any cell in the tissue is on average 3.3. Only 31% of model-generated tissues have a maximum cell-fate potential of 5 cell-types. When instead we also include cell-types that can be produced through indirect differentiation, the maximum cell-fate potential averaged across tissues is 3.7, and 57% of model-generated tissues have a maximum cell-fate potential of 5 cell-types. In FigS.3, we describe how maximum cell-fate potential depends on model parameters: Similarly to average cell-fate potential, maximum cell-fate potential (direct differentiation) increases with parameters $P_{den}$ and $P_{diff}$ (Pearson's correlation coefficients: 0.14, 0.70, respectively) and decreases with $P_{stable}$ (Pearson's correlation coefficient: -0.16). The corresponding Pearson's correlation coefficients with model parameters for maximum cell-fate potential (indirect differentiation incl.) are: 0.14, 0.66 and -0.16 with parameters $P_{den}$, $P_{diff}$ and $P_{stable}$, respectively.

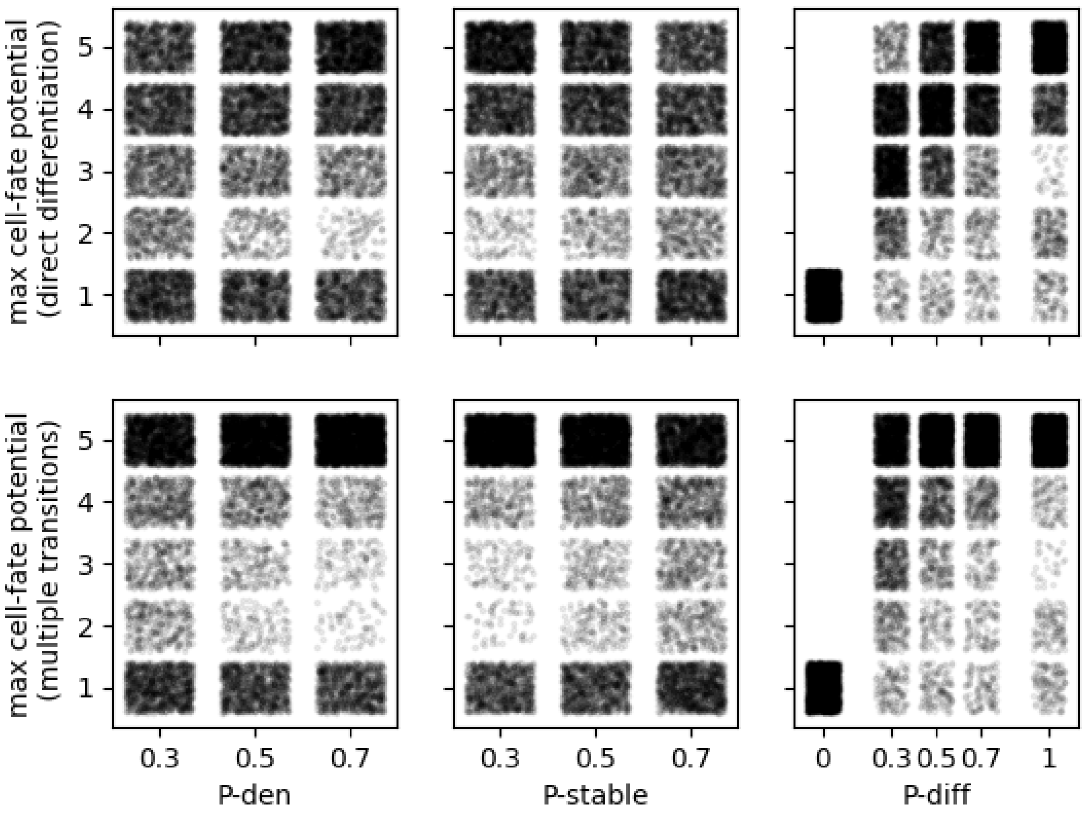


FigS. 3: Scatter-plots showing the dependence on model parameters of maximum cell-fate potential of tissue cell-types, where (Row-1) cell-fate only counts cell-types that can be produced through direct differentiation, (Row-2) cell-fate also considers indirect mapping through multiple differentiation steps. From left to right, the x-axis represents $P_{den}$, $P_{stable}$, $P_{diff}$. Each point represents a model-generated tissue. Noise has been added to the position of points to make density more apparent.

### 3 Tissue spatial properties

We characterize spatial properties of model-generated tissues using 5 properties (FigS.4):

1. fullness: tissue fullness increases strongly with parameter $F_{adj}$ (Pearson's correlation coefficient: 0.65). It decreases with $P_{stable}$ and $P_{diff}$, parameters which preclude migration (Pearson's correlation coefficient: -0.10, -0.27, respectively), and increases modestly with $P_{den}$ (Pearson's correlation coefficient: 0.10).

2. number of unique domains: Number of unique tissue domains decreases strongly with parameter $P_{diff}$ (Pearson's correlation coefficient: -0.47). It decreases to a modest extent with $P_{stable}$ and $F_{adj}$ (Pearson's correlation coefficient: -0.13, -0.15, respectively), and increases modestly with $P_{den}$ (Pearson's correlation coefficient: 0.14).

3. coverage equality: This property represents the relative proportion of the tissue covered by different tissue domains. Coverage equality increases strongly with parameter $F_{adj}$ (Pearson's correlation coefficient: 0.39). It depends very weakly on values of the other parameters $P_{den}$, $P_{stable}$ and $P_{diff}$ (Pearson's correlation coefficient: 0.08, -0.08, -0.10 respectively).

4. dispersity: Average dispersity across all domains within a tissue tends to increase with parameters $P_{den}$ (Pearson's correlation coefficient: 0.24). It decreases with parameters $F_{adj}$ $P_{stable}$ and $P_{diff}$ (Pearson's correlation coefficient: -0.09, -0.25, -0.34 respectively).

5. mixedness: Average mixedness across all domains within a tissue displays a similar dependence as average dispersity: It tends to increase with parameters $P_{den}$ (Pearson's correlation coefficient: 0.23). It decreases with parameters $F_{adj}$ $P_{stable}$ and $P_{diff}$ (Pearson's correlation coefficient: -0.24, -0.22, -0.44 respectively).

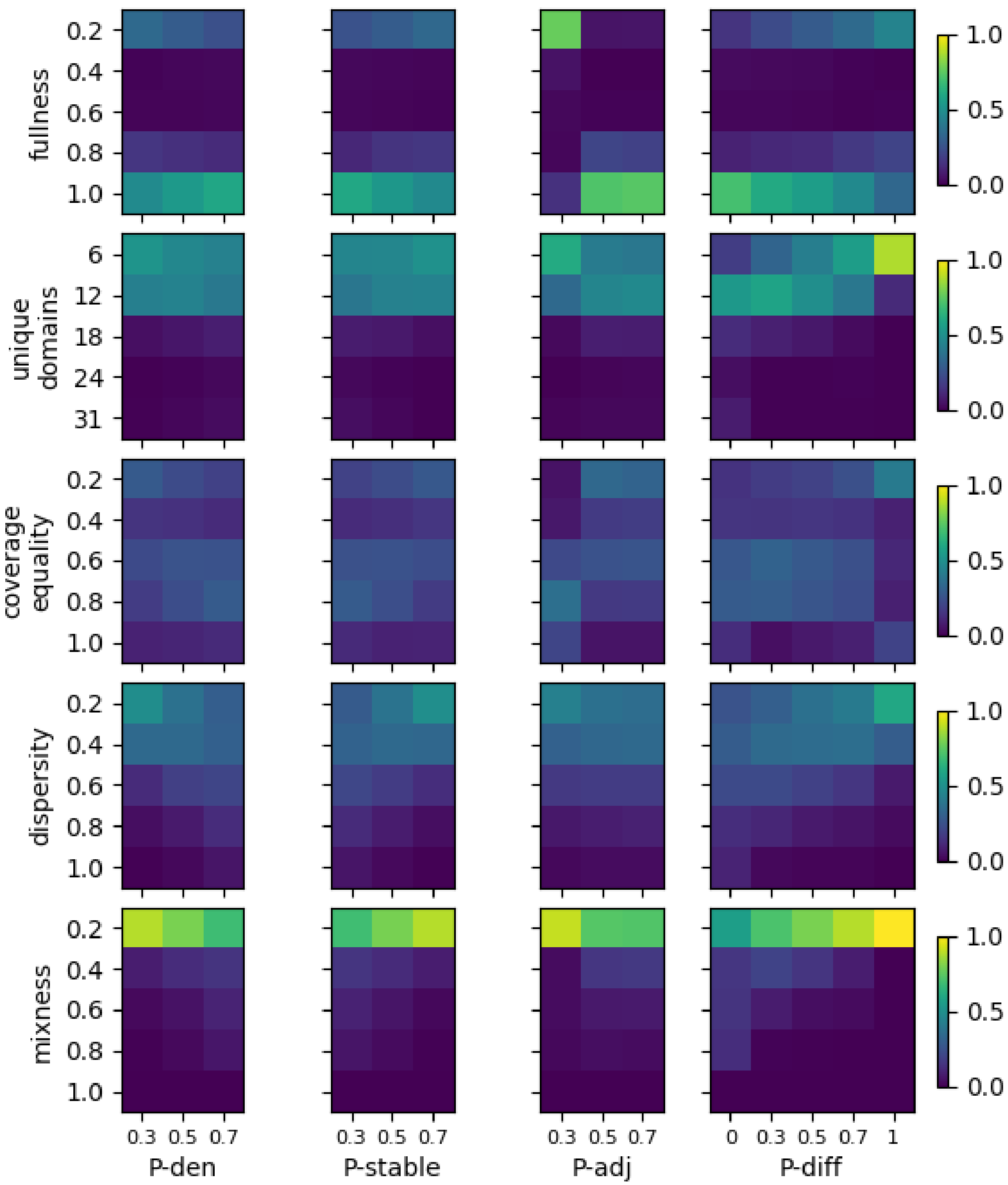


FigS. 4: Heatmaps displaying dependence on model parameters of the 5 spatial properties using which we characterize model-generated tissues: fullness (row-1), number of unique domains (row-2), coverage equality (row-3), dispersity (row-4) and mixedness (row-5). In each heatmap, the rows represent binned values of the corresponding spatial properties and the columns represent values of model parameters: $P_{den}$ (column-1), $P_{stable}$ (column-2), $F_{adj}$ (column-3) and $P_{diff}$ (column-4). Each element (i,j) of each heatmap represents the fraction of tissues generated with parameter value corresponding to the $j^{th}$ column that have a value of spatial property corresponding to the $i^{th}$ row. Colorbars describing colors for heatmaps belonging to different rows are provided on the right.

We then perform a Principal Components Analysis to reduce the dimensionality of the space of tissue spatial properties (FigS.5). In FigS.6, we show how the two significant Principal Components (PC1 and PC2) depend on model parameters:

1. PC1: PC1 displays a similar dependence as average dispersity: It tends to increase with parameters $P_{den}$ (Pearson's correlation

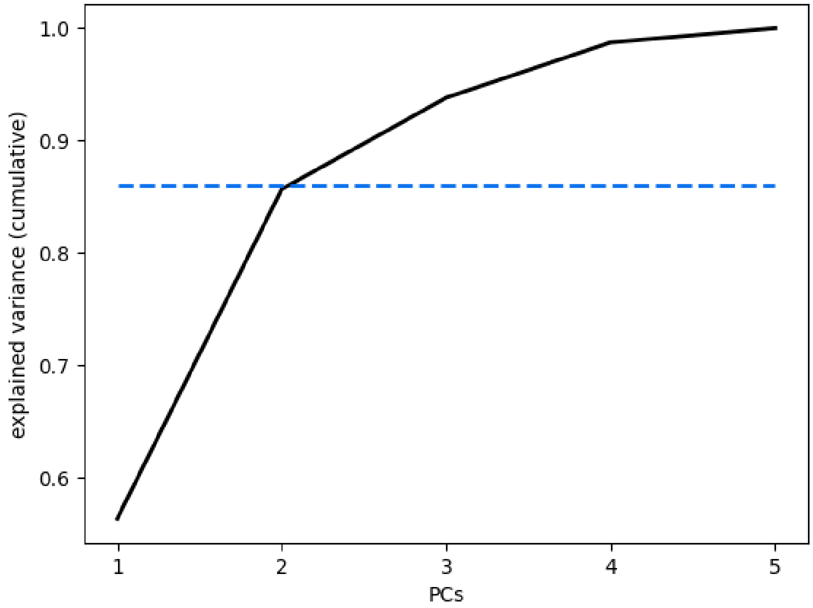


FigS. 5: Elbow plot showing the variance in the data explained by each PC. PCs 1 and 2 explain 86% of the variance in the data as indicated by the dotted blue line.

coefficient: 0.22). It decreases with parameters $F_{adj}$ $P_{stable}$ and $P_{diff}$ (Pearson's correlation coefficient: -0.24, -0.21, -0.46 respectively).

2. PC2: PC2 displays a similar dependence as tissue fullness: It increases strongly with parameter $F_{adj}$ (Pearson's correlation coefficient: 0.57). PC2 decreases modelstly with $P_{stable}$ (Pearson's correlation coefficient: -0.02), and increases modestly with $P_{den}$ and $P_{diff}$ (Pearson's correlation coefficient: 0.02, 0.06, respectively).

### 4 Clusters of tissue-types versus sectors

We subdivide the space of tissue configurations defined by PC1 and PC2 into 4 sectors: *stunted, sparse-contiguous, full contiguous and random-disperse*. When we performed hierarchical clustering of model generated tissues based on their 5 spatial properties (fullness, number of unique domains, coverage equality, mixedness, dispersity), tissues separated into 3 robust clusters (FigS.7(A)). In FigS.7(B), we compare sector boundaries against cluster assignments: essentially, the sectors *stunted* and *sparse-contiguous* represent a finer subdivision of tissues belonging to the same cluster.

### 5 Tissue developmental trajectories

On average, developmental trajectories of model-generated tissues involve 5.7 sector-crossings (FigS.8, Column-1). While trajectories of tissues that end up in the stunted and sparse-contiguous sectors involve very few sector-crossings (on average, 0.5 crossings and 2.4 crossings, respectively), trajectories of tissues that end up in full-contiguous and random-disperse sectors involve many more sector crossings (on average, 7.2 crossings and 9.5 crossings, respectively).

The number of time-steps within which tissues reach their final sectors is on average 41 time-steps (FigS.8, Column-2). While trajectories of tissues that end up in the stunted and sparse-contiguous sectors reach the final sector within very few time-steps (on average, 0.71 and 13, respectively), trajectories of tissues that end up in full-contiguous and random-disperse sectors take many more time-steps (on average, 61 and 51, respectively).

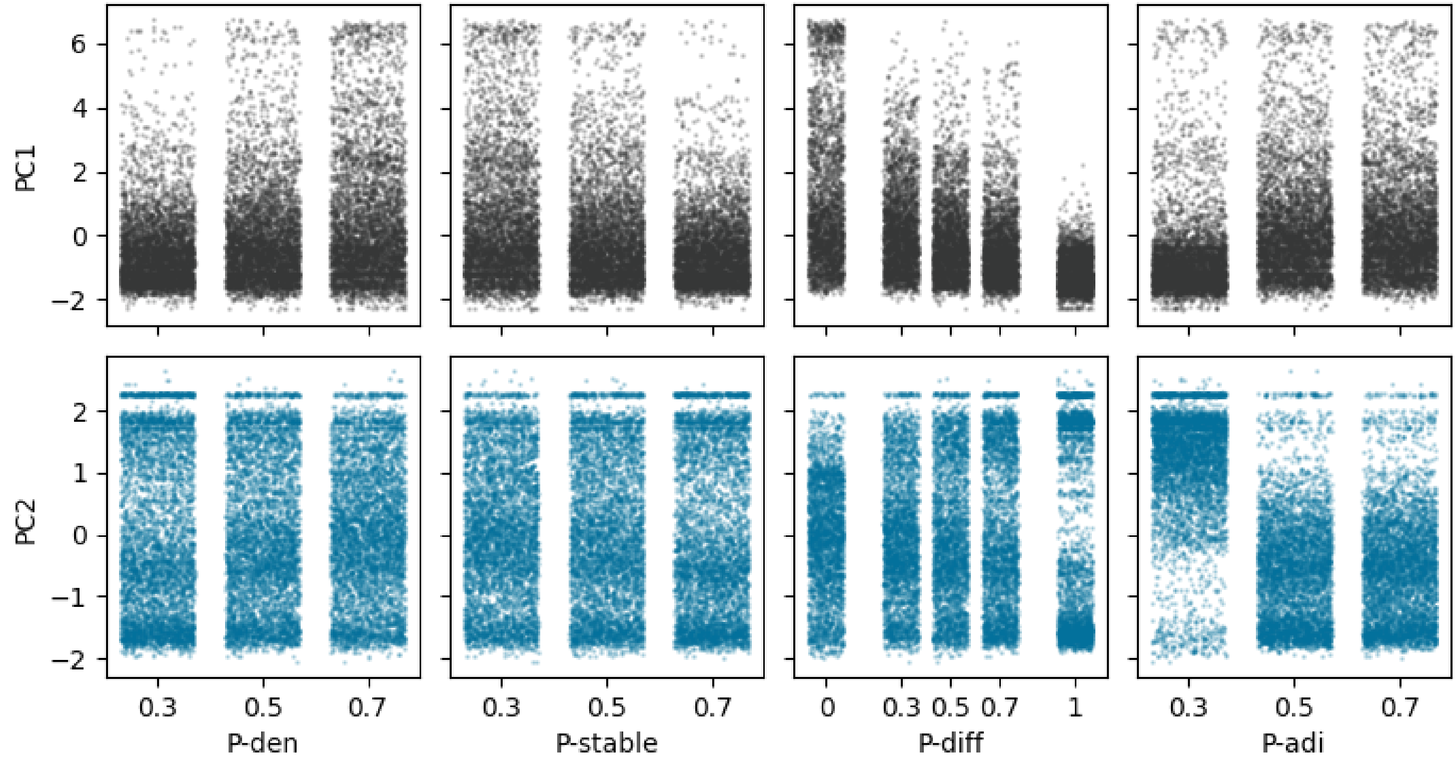


FigS. 6: Scatter-plots showing the dependence on model parameters of Principal Components (Row-1) PC1 and (Row-2) PC2 which represent tissue spatial properties. From left to right, the x-axis represents $P_{den}$, $P_{stable}$, $P_{diff}$. Each point represents a model-generated tissue. Noise has been added to the position of points to make density more apparent.

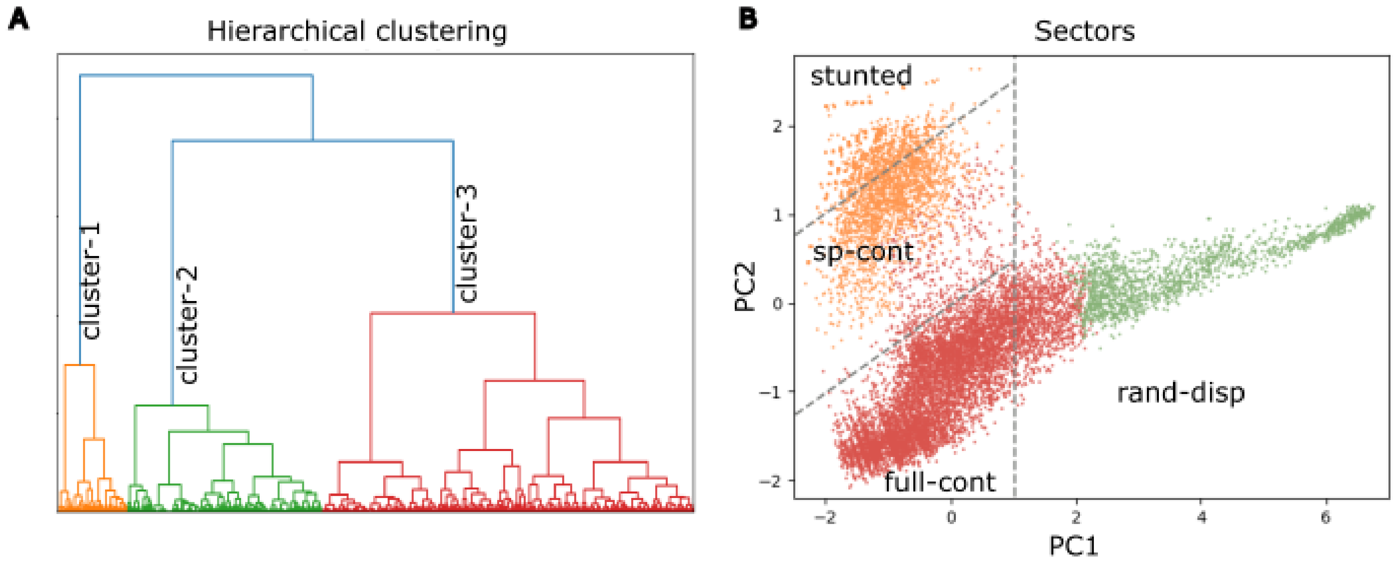


FigS. 7: Hierarchical clustering of model generated tissues. (A) Clustering of tissues based on 5 spatial properties yields 3 clusters (cluster-1 is orange, cluster-2 is green, cluster-3 is red). (B) Comparison of tissue clusters against tissue sectors. Each dot represents a distinct tissue. Dot color represents its cluster assignment. Dotted grey lines represent sector boundaries. Each sector is labeled as stunted, sp-cont, full-cont and rand-disp.

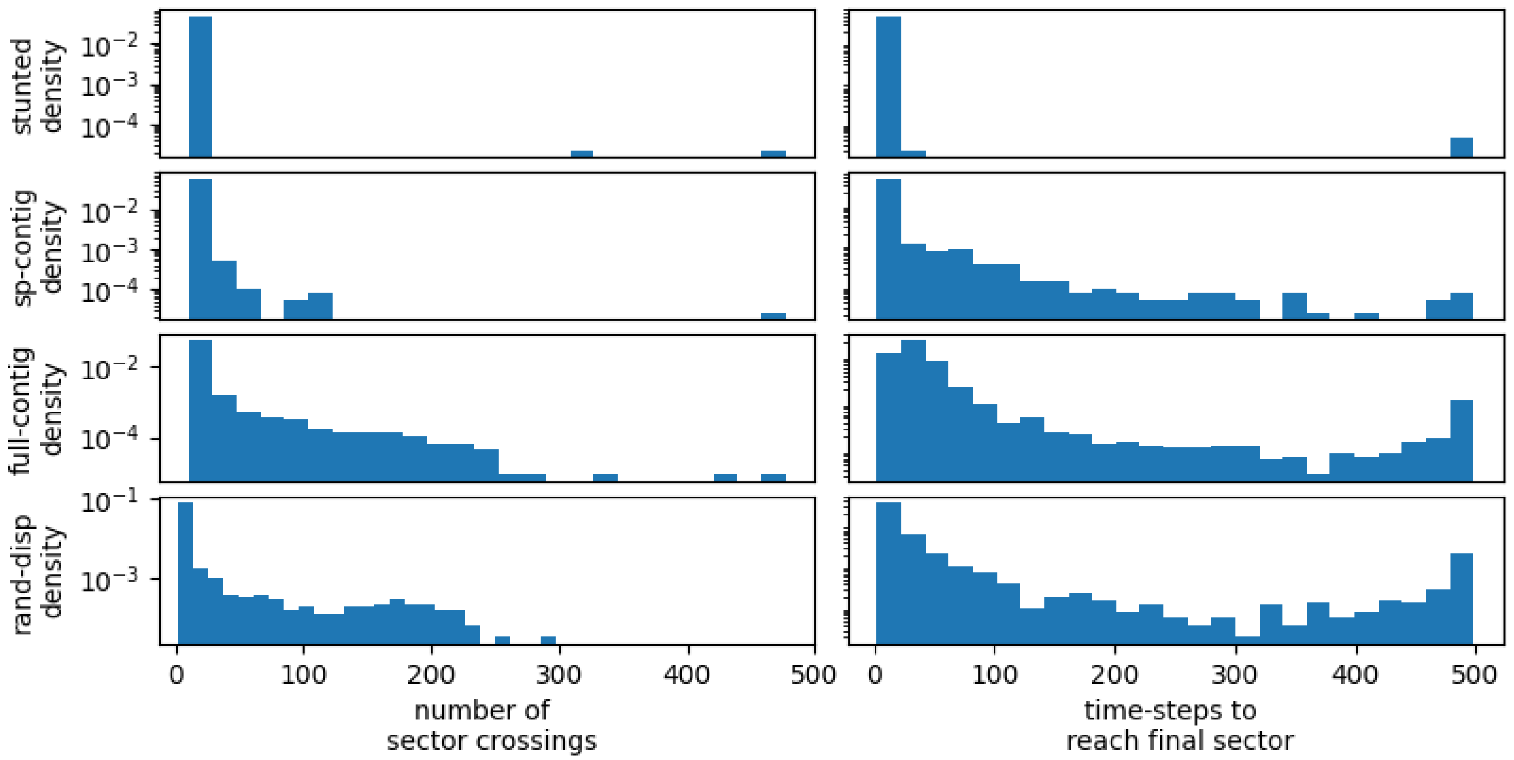


FigS. 8: Histograms for (Column-1) the number of sector crossings, (Column-2) number of time-steps to reach the final sector. Each row represents tissues in different sectors: (Row-1) stunted, (Row-2) sparse-contiguous, (Row-3) full-contiguous and (Row-4) random-disperse.

### 6 Tissue healing from typeB injuries

For each tissue, we perform 25 independent injuries, where we ablate a single randomly chosen cell-type from all grid-squares adjacent to a randomly chosen grid-square. We analyse how this tissue responds to this injury by counting the number of times out of 25 that it recovers, doesn't recover, or degrades further. Almost all tissues which are able to recover (recovery-fraction¿0.5) are in the full-contiguous sector (FigS.9). We describe in FigS.10 how tissue recovery depends on model parameters:

1. recovery-fraction: The fraction of times out of 25 that a tissue recovers increases with parameter $P_{stable}$, $F_{adj}$ and $P_{diff}$ (Pearson's correlation coefficient: 0.13, 0.44, 0.24 respectively). It decreases with $P_{den}$ (Pearson's correlation coefficient: -0.11).

2. no-recovery-fraction: The fraction of times out of 25 that a tissue neither recovers, nor gets worse increases with parameter $P_{stable}$ and $P_{diff}$ (Pearson's correlation coefficient: 0.10, 0.09 respectively). It decreases with $P_{den}$, $F_{adj}$ (Pearson's correlation coefficient: -0.09, -0.48, respectively).

3. degradation-fraction: The fraction of times out of 25 that a tissue neither recovers, nor gets worse increases with parameter $F_{adj}$, $P_{den}$ (Pearson's correlation coefficient: 0.01, 0.15 respectively). It decreases with $P_{diff}$, $P_{stable}$ (Pearson's correlation coefficient: -0.27, -0.18, respectively).

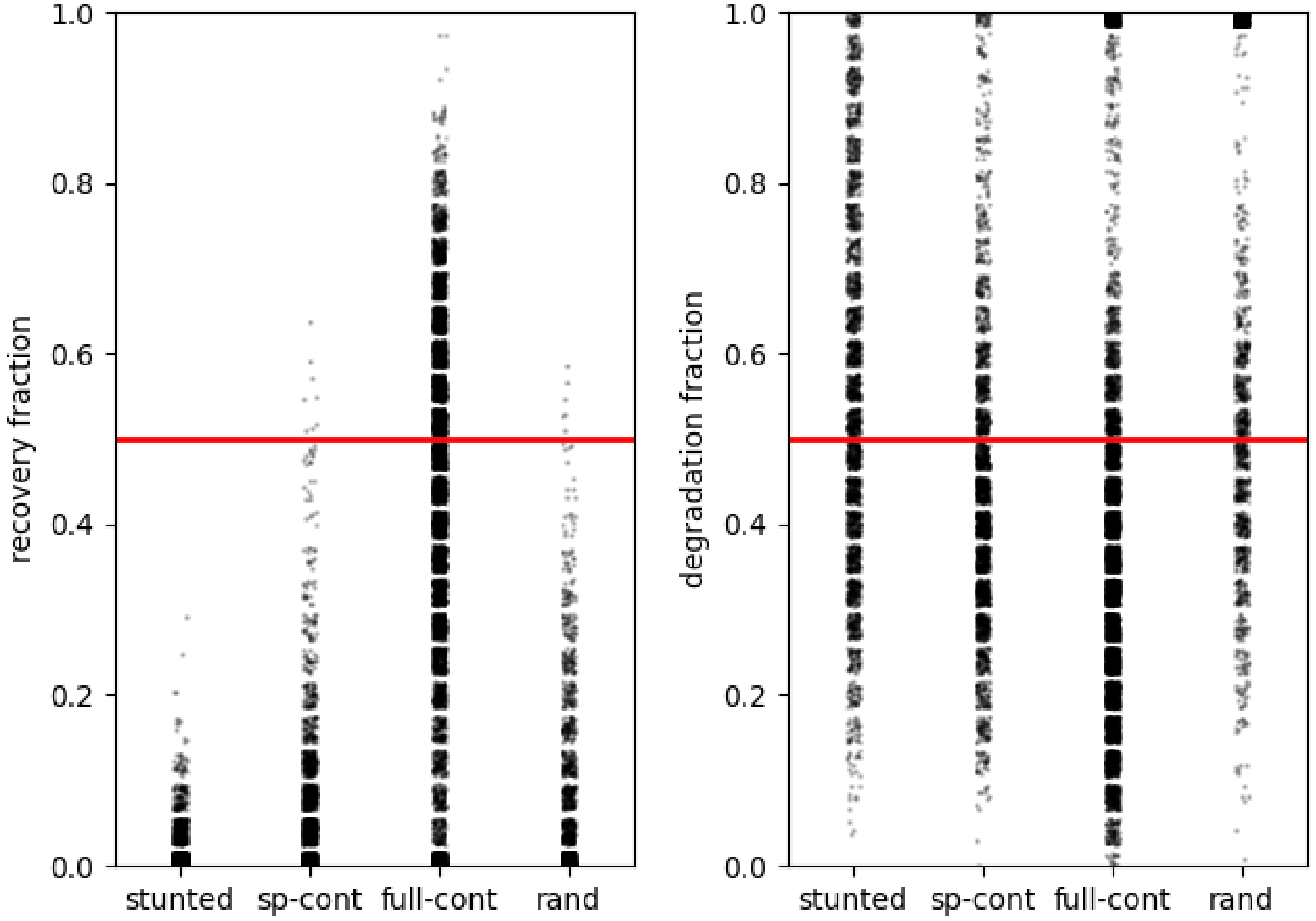


FigS. 9: Scatterplots displaying the extent of tissue healing for tissues (tissue types given by the x-axes) in response to typeB injuries. Each point represents a distinct tissue. Red lines indicate a recovery fraction/ degradation fraction of 0.5. (Left plot) y-axis represents binned values for recovery-fraction, (Right plot) y-axis represents binned values for degradation-fraction

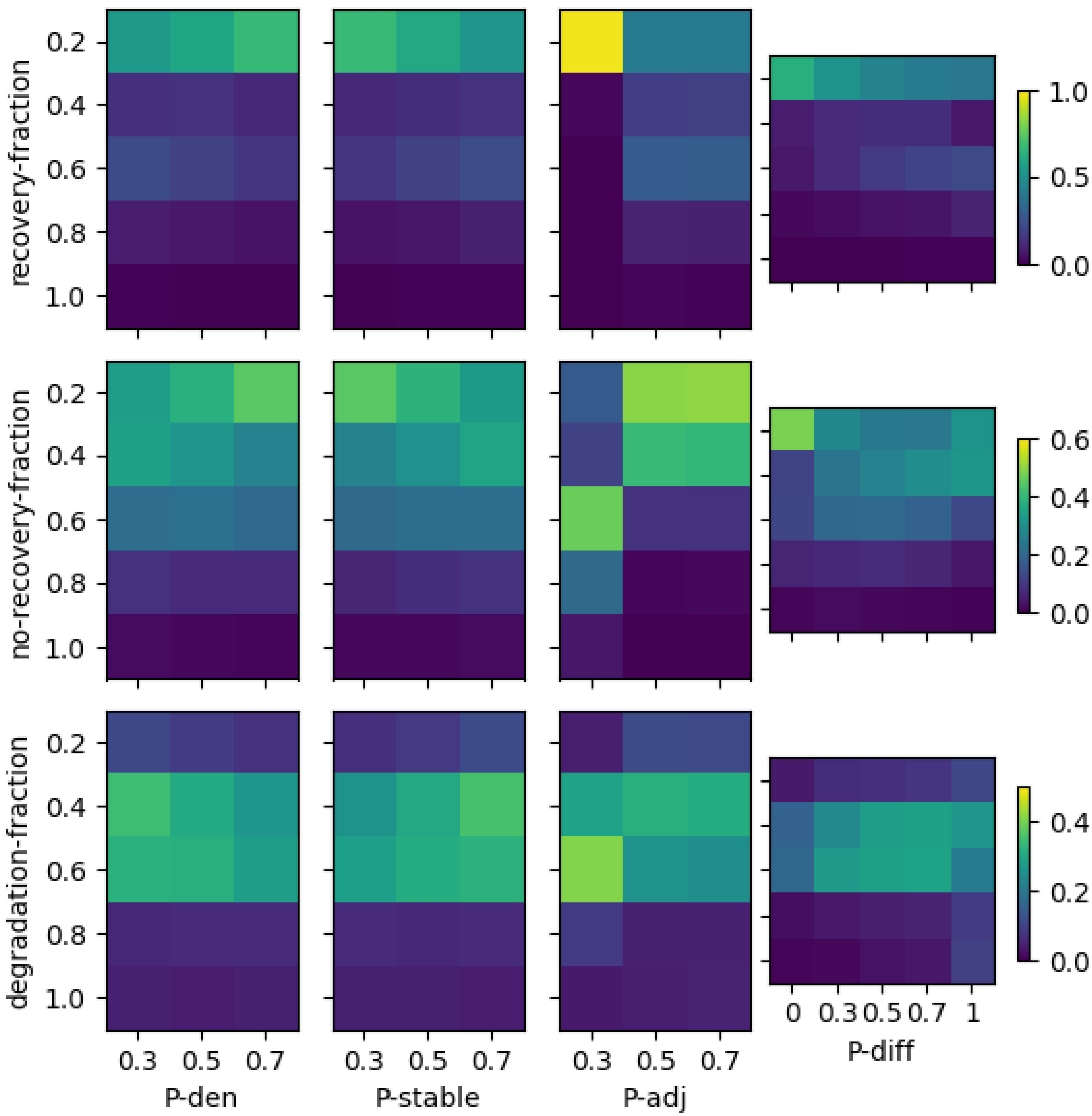


FigS. 10: Heatmaps displaying dependence of tissue healing in response to typeB injuries on model parameters. Rows represent binned values of (Row-1) recovery-fraction, (Row-2) no-recovery-fraction, (Row-3) degradation fraction, and columns represent model parameters: (Column-1) $P_{den}$, (Column-2) $P_{stable}$, (Column-3) $F_{adj}$ and (Column-4) $P_{diff}$. Each element (i,j) of each heatmap represents the number of tissues with a value of (Row-1) recovery-fraction, (Row-2) no-recovery-fraction, (Row-3) degradation-fraction corresponding to the $i^{th}$ row, and parameter value corresponding to the $j^{th}$ column. Colorbars describing colors for heatmaps are provided on the right.
